# An ancient receptor family illuminates the evolution of animal sensation

**DOI:** 10.64898/2026.06.15.732144

**Authors:** Maxwell C. Coyle, John K. Nunnally, Yeun-Hyeok Shin, Yury A. Trofimov, Irina I. Veretenenko, Jack Thibodeau, Irina A. Talyzina, Riya Sivakumar, Roman G. Efremov, Jon Clardy, Alexander I. Sobolevsky, Nicholas W. Bellono

## Abstract

Animals use nervous systems to sense and respond to their environment. Yet single-celled organisms can also detect cues to execute diverse behaviors, suggesting that core components for animal sensation predate multicellularity and nervous systems. Here, we report that choanoflagellates, the closest living animal relatives, use an ancient sensory receptor family to detect bacterial prey. These receptors are related to transient receptor potential ion channels but are distinguished by WD40 domains, defining TRPW. TRPW1 detects specific bacterial lipids to modulate flagellar beating, providing a mechanism for attraction towards prey. TRPW emerged in early eukaryotes and reveals ancestral architectural and ligand-binding features that predate animal somatosensory receptors. In multicellular choanoflagellates, TRPW1 elicits collective responses, linking bacterial ecology to the evolution of receptors, sensory organelles, and multicellular life.

## Introduction

Animals use sensory receptors, specialized cell types, and multi-cellular neural circuits to sense and respond to their environment. Yet, animals evolved from unicellular eukaryotes (protists) that can execute complex behaviors even as single cells (1–3). Less is known about how protists sense and respond to their environment or how sensory systems co-evolve alongside transitions from unicellular to multicellular life. Choanoflagellates (choanos) are well-positioned to provide insights into these questions because they are the closest living relatives of animals and can transition flexibly between unicellular and multicellular states (4, 5). Here, we use choanoflagellates to uncover and investigate a novel ion channel receptor family with deep eukaryotic ancestry that is closely related to, and predates, animal somatosensory receptors. We show that one of these receptors detects specific lipids from prey bacteria, driving flagellar responses and multicellular coordination, revealing how bacterial ecology shapes the evolution of receptors, sensory organelles, and multicellular processing systems.

## Results

### An ion channel receptor family for prey bacteria in choanoflagellates

Choanos are obligate bacterivores that must locate and consume specific prey bacteria (*6, 7*). Furthermore, the choano species *Salpingoeca rosetta* (*S. rosetta*) responds to sulfonolipids from the prey bacterium *Algoriphagus machipongonensis (A. mac*) by forming multicellular rosettes (*8, 9*). Thus, we reasoned that choanos may detect specific molecules to guide them towards prey. Indeed, when we exposed *S. rosetta* to *A. mac* biofilms, we found that choanos were attracted to and swarmed near the biofilm surface, suggesting the detection of local molecules (Figure 1A, Movie S1). How do choanos sense and respond to their environment? Chemoreceptors exhibit a dynamic history of lineage-specific expansions and contractions that correlate with the disparate complexity of chemical ecology (*10*). Guided by this rationale, we sought to identify gene expansions that could encode chemoreceptors for bacterial molecules in choanos. Using OrthoFinder, we identified an expanded gene family with 17 members in *S. rosetta* that exhibited a distinct architecture: a voltage-gated ion channel superfamily transmembrane domain and variable numbers of N-terminal WD40 repeats. Phylogenetic analysis of the transmembrane domains showed that this family is most closely related to type I transient receptor potential (TRP) channels including TRPA, TRPV, and TRPM, which serve diverse sensory functions in animals (*11*) (Figure 1B, Figure S1A-E). However, these genes constitute their own family, as indicated by their phylogenetic grouping with one another and the presence of WD40 domains, which are not found in other TRP families. Furthermore, the distinctive gene architecture of these choano receptors is found across diverse eukaryotes, pointing to a deep ancestry followed by secondary loss in animals (Figure 1C, Materials and Methods). Considering their distinct phylogenetic distribution and unique WD40 domain architecture, we herein refer to this family as TRPW.

**Figure 1:**
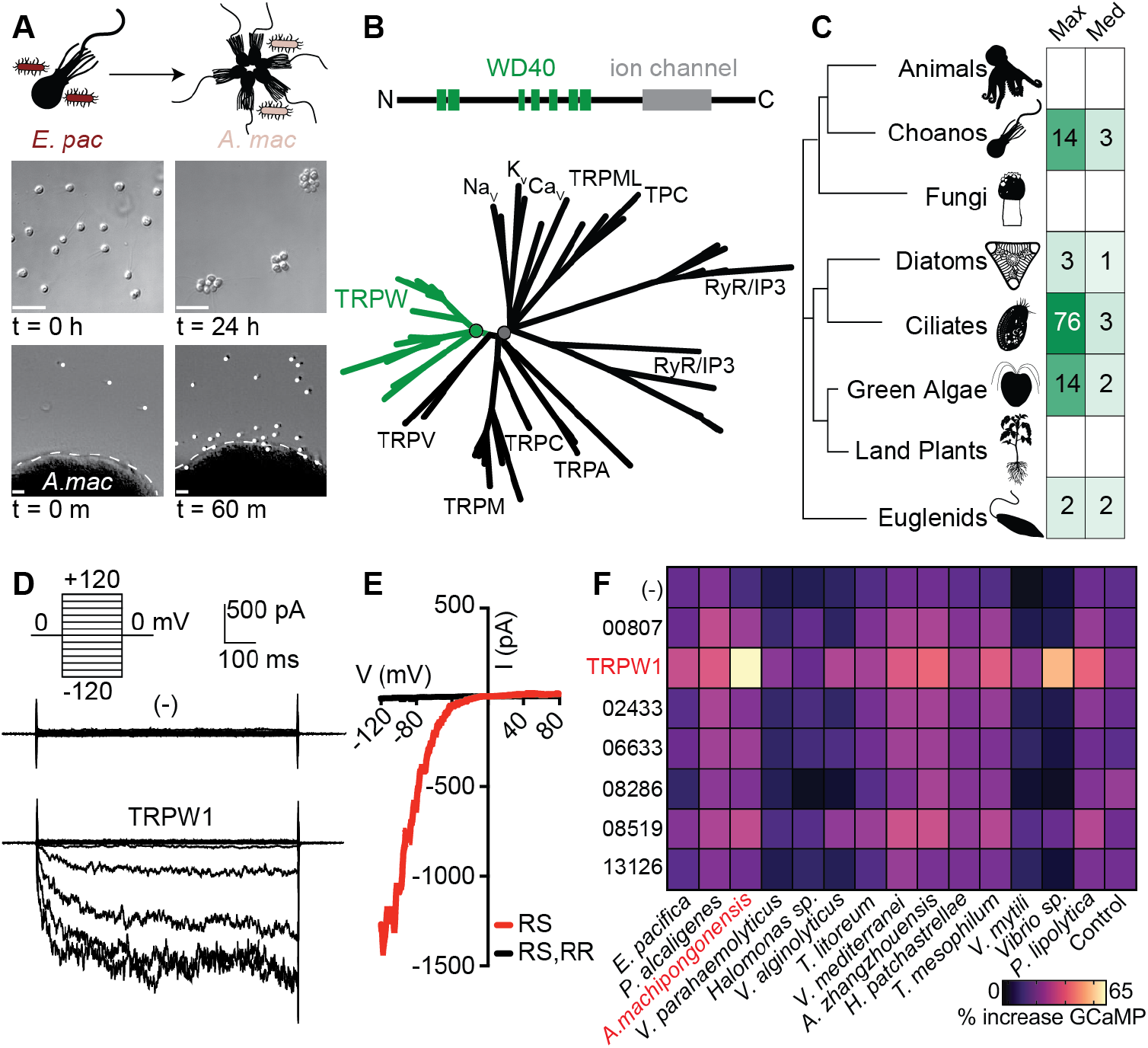
An ion channel receptor family in choanoflagellates. **(A)** Choanoflagellate (choano, *Salpingoeca rosetta*) attraction behavior and multicellular rosettes are induced by *Algoriphagus machipongonensis (A. mac*), a prey bacterium co-isolated from the same natural habitat, a marsh in Hog Island, Virginia. Rosette formation does not occur when *S. rosetta* is fed with another bacterium, *Echinicola pacifica* (*E. pac*) (*8*). Scale bars = 20 μm. See Movie S1. **(B)** *S. rosetta* encodes 17 members of an expanded gene family containing a voltage-gated ion channel superfamily transmembrane domain and WD40 repeats. These genes form a highly supported clade (91/94 support by UFboot/SH-aLRT, green node) related to type I TRP channels (82/92 support, gray node). Phylogenies from diverse animals and choanos in Figure S1D. **(C)** The TRPW domain architecture is found across diverse eukaryotic lineages. See Materials and Methods for discussion of two false positive hits in animals. **(D)** TRPW1 expression in HEK293T cells mediates an inwardly rectifying current in response to synthetic agonist RS 39604 (RS, 100 μM) identified in small molecule screens (Fig. S1G). **(E)** RS-evoked TRPW1 currents were inhibited by 10 μM ruthenium red (RR). Representative of n = 8 cells. **(F)** Among *S. rosett»*RPW genes (labeled by gene accession number), TRPW1 responded most strongly to an extract from the co-isolated bacterium *A. mac*, compared to extracts from 13 other marine bacterial species. Activity is percentage increase in GCaMP. Heatmap indicates average of n = 3 Ca^2+^ imaging replicates.

We next asked whether TRPWs encode ligand-activated ion channels that could underlie detection of bacteria. We cloned and heterologously expressed the most highly transcribed TRPW genes in HEK293 cells to test responses to a 2000-molecule library. We found specific agonists, including RS 39604 (RS, a highly substituted piperidine-based molecule) that activates one of the TRPW proteins, which we call TRPW1. (Figure S1G-J). With RS and other ligands, we used patch-clamp electrophysiology to demonstrate that TRPW1 is indeed a ligand-activated, calcium-permeable, nonselective cation channel that was inhibited by ruthenium red (Figure 1D,E, Figure S1K-N). To ask if TRPWs are activated by ecologically and behaviorally relevant molecules, we tested extracts from 14 different cultured marine bacteria against 7 heterologously expressed TRPWs.

Remarkably, TRPW1 was most strongly activated by extracts from *A. mac*, the rosette-inducing bacterial species that was co-isolated with *S. rosetta* (*8*) (Figure 1F).

How do specific prey bacteria activate TRPW1? Gram negative bacteria like *A. mac* release outer membrane vesicles (OMVs) that can contain signaling lipids (9, 12). Indeed, OMVs strongly activated TRPW1 even after exposure to boiling temperatures that denature peptides, suggesting a lipid could constitute the active biomolecule (Figure 2A). To isolate active molecules from *A. mac*, we used iterative, activity-guided, high-performance liquid chromatography coupled to mass spectrometry (HPLC-MS) and nuclear magnetic resonance (NMR) to identify a mono-unsaturated branched chain fatty acid, which we refer to as 15M-9E (Figure 2B,C, Figure S2A-I). Purified 15M-9E evoked dose-dependent TRPW1 activity with similar voltage-dependence as synthetic agonists (Figure 2D,E). Consistent with the high specificity of bioactivity elicited by this bacterial strain, the fully saturated version of this lipid differs by only two hydrogen atoms but did not activate TRPW1 (Figure 2E).

**Figure 2:**
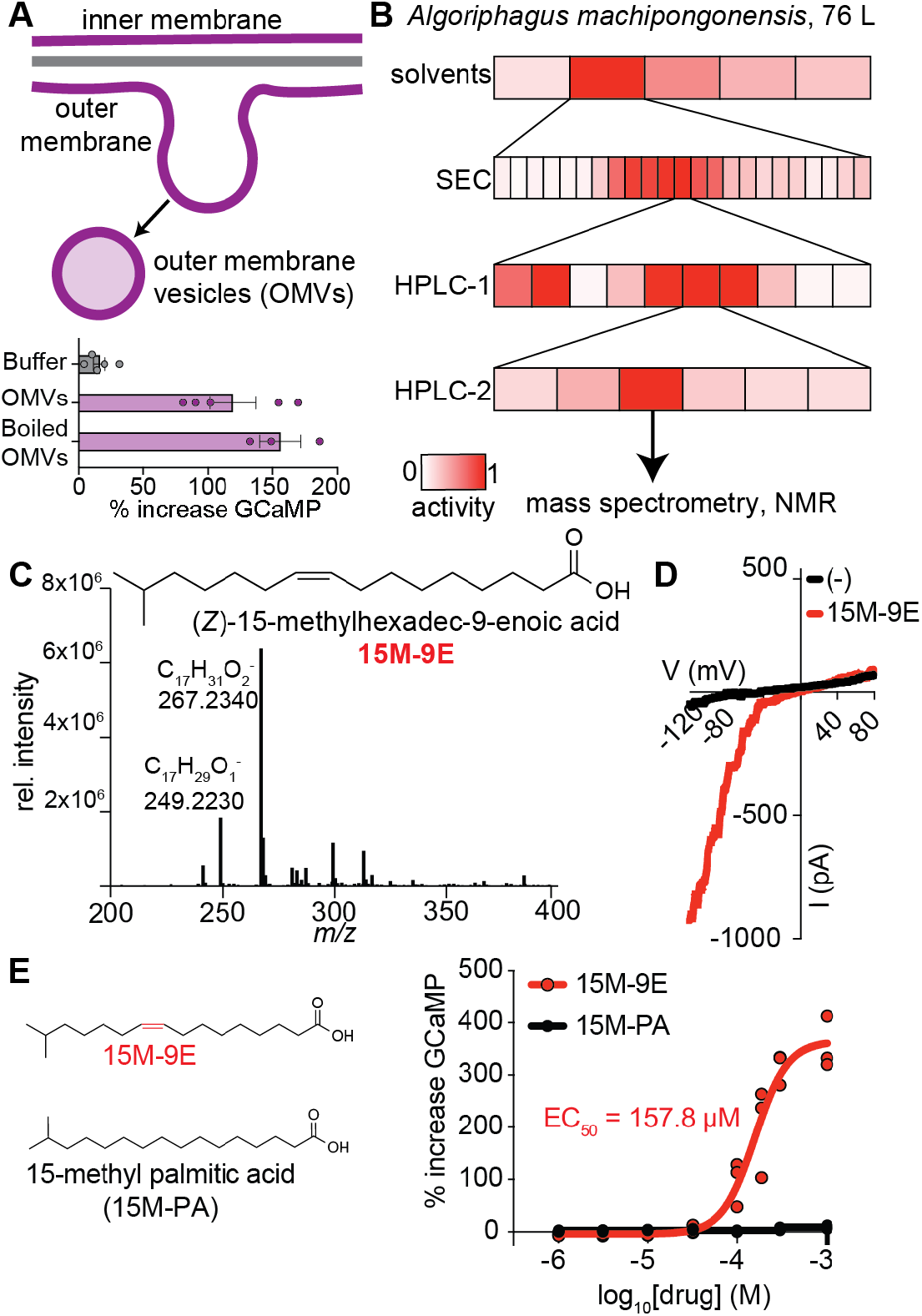
TRPW1 detects specific bacterial prey lipids. **(A)** *A. mac* gram-negative bacteria contain inner and outer membranes. *A. mac* releases outer membrane vesicles that activated TRPW1 even when boiled to denature proteins. Each dot indicates average across a biological replicate. Error bars show S.E.M. p = 0.0034 OMV vs. control, p = 0.0081 boiled OMV vs. control, unpaired t-test. **(B**) TRPW1 activity-guided fractionation of *A. mac* cultures identified a pure fraction for small molecule identification. Activity normalized to the most active fraction. See Fig. S2A-D. **(C)** Mass spectrometry and NMR (Fig. S2E-I) identifies the chemical structure of the *A. mac* TRPW1 agonist as 15M-9E. The two most abundant *m/z* peaks correspond to the negative ion of the branched chain fatty acid and a dehydration product. **(D)** Current-voltage relationship of TRPW1 activity in response to 187 μM 15M-9E. Representative of n = 5 cells. **(E)** 15M-9E activates TRPW1, but 15-methyl palmitic acid (15M-PA) is inactive to 1 mM despite differing in only one carbon-carbon double bond. 15M-9E EC_50_ = 157.8 μM (95% CI from 126.4 μM to 195.7 μM). n = 3 Ca^2+^ imaging replicates.

### TRPW1 regulates flagellar activity to promote attraction to agonists

Considering the specific activation of TRPW1 by an *A. mac* lipid, we wondered if TRPW1 ligands could recapitulate choano attraction near prey (Figure 1A). To test this hypothesis, we embedded small agar pucks with RS (a more soluble agonist than 15M-9E) to model diffusion of bioactive molecules within OMVs away from surfaces. This resulted in significant cell attraction to the puck compared with control, thereby suggesting a molecular basis for chemosensory behavior in *S. rosetta* (Figure 3A). Because this attraction appeared to be due to slower swimming behavior at higher concentrations of agonist near the puck surface, we next exposed cells to constant concentrations of agonists. We found that *S. rosetta* cells exposed to 15M-9E periodically stopped swimming, manifesting in a slower average speed compared to vehicle control (Figure 3B). To ask whether TRPW1 is responsible for this phenotype, we used CRISPR-Cas9 genome editing to generate a TRPW1^KO^ strain, which exhibited a partially altered swimming phenotype (Figure 3B, Figure S3A-B). Importantly, TRPW1^KO^ did not show any impairment in growth, phagocytosis, or rosette formation (Figure S3C-E).

**Figure 3:**
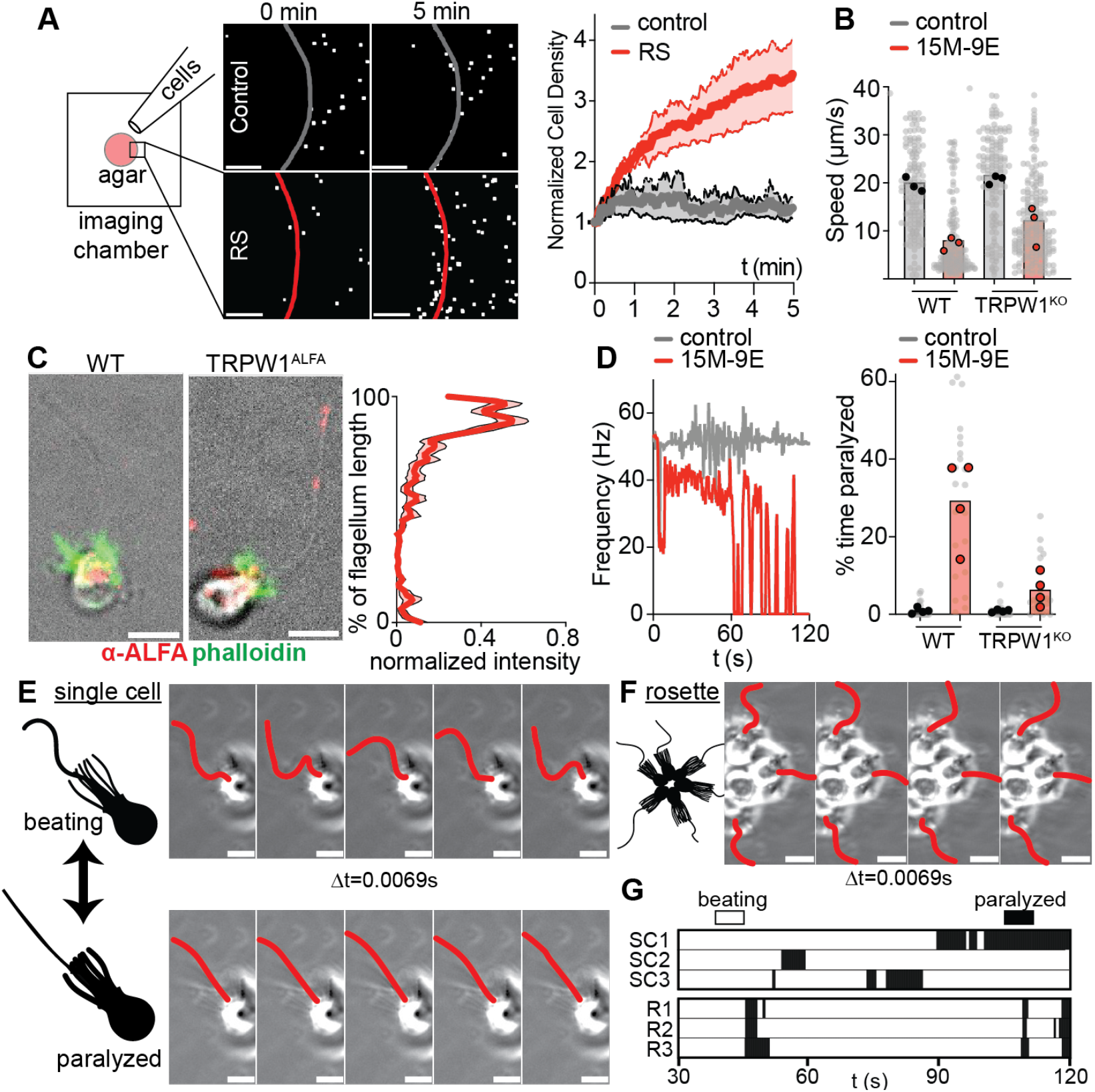
TRPW1 regulates flagellar activity and attraction to agonists. **(A)** *S. rosetta* cells were attracted to agar pucks containing TRPW1 agonist (RS, 500 μM) but not vehicle control (DMSO). n = 3 experiments, lines indicate average and S.E.M. for cell density in a field of view at the edge of the agar puck. Representative views shown for 0 and 5 min. Scale bar = 200 μm. **(B)** 15M-9E led to slower average swimming speed. Gray dots show individual tracks, large dots weighted averages for biological replicates. p = 0.0005 for WT control vs 15M-9E, t-test; for 15M-9E treatment, genotype x treatment interaction for means was n.s., but distinct distributions were supported for WT vs. TRPW1^KO^ (coefficient of variation, p = 0.0175, t-test). **(C)** TRPW1^ALFA^ localization at the distal end of the choanoflagellate flagellum. Phalloidin staining indicates the microvillar collar. Scale bar = 5 μm. Average intensity trace profile with S.E.M. displayed as bounding lines for n = 20 flagella. **(D)** 75 μM 15M-9E caused sporadic flagellar paralysis, which was reduced in choanos with TRPW1 genetic deletion (TRPW1^KO^). p = 0.0001 for drug treatment, p = 0.0023 for genotype x treatment interaction, two-way ANOVA. Drug addition occurs during first 5 seconds. **(E)** 75 μM 15M-9E induced unicellular *S. rosetta* to sporadically switch between a state of actively beating flagella and a state of flagellar paralysis. Scale bar = 5 μm. **(F)** Multicellular *S. rosetta* rosettes show coordinated flagellar paralysis in respond to TRPW1 ligands (50 μM RS). Scale bar = 5 μm. Percentage of coordinated first paralysis events = 65.7 +/-15.0 % S.D. across n = 3 biological replicates. **(G)** Time-course data showing transitions between active beating and paralysis for 3 independent single cells (SC1-3) and 3 cells connected as part of a rosette (R1-3). Time is seconds after drug addition, with no responses observed in first 30 s.

To understand how TRPW1 regulates behavior, we localized TRPW1 by using immunofluorescence together with a genetically inserted C-terminal ALFA peptide (*13*) (TRPW1^ALFA^, Figure S3A,F-G). TRPW1 local-ized to the distal end of the flagellum, in agreement with the regulation of TRPW expression as part of the core flagellar transcriptional program (*14*) (Figure 3C, Figure S3H). Flagellar localization of choano sensory receptors lends further support to hypotheses that cilia and flagella evolved sensory functions deep in eukaryotic ancestry (*15–17*). Considering TRPW1’s flagellar localization, we next asked whether TRPW1 activation alters flagellar activity to control swimming behavior. Using a microfluidic chamber to deliver 15M-9E and a high-speed camer»o record flagellar activity, we found that TRPW1 agonists caused cells to switch between an actively beating and a paralyzed state, compared to continuous flagellar beating in control conditions (Figure 3D,E). Intriguingly, calcium influx induces ciliary paralysis across diverse animal cells, and signal transduction components are conserved in choanos and animals (18, 19). Finally, 15M-9E failed to induce robust flagellar paralysis in TRPW1^KO^, demonstrating the importance of TRPW1 for detecting 15M-9E to control flagellar beating (Figure 3D). Thus, bacterial lipids activate TRPW1 to regulate flagellar activity and swimming, demonstrating one mechanism for attraction to prey sources.

Choanos can form multicellular rosettes that are connected by intercellular bridges (20) and appear to be electrically coupled (21), raising the possibility that sensory stimuli could elicit coordinated multicellular responses, similar to those observed in nervous systems (22). In response to TRPW1 agonists, both single cells and rosettes stochastically switch between paralyzed and beating states (Figure 3E,F). Strikingly, cells in rosettes repeatedly arrested simultaneously in response to agonists. These arrest events did not occur immediately upon exposure, but stochastically in a time window during continuous exposure, demonstrating coordinated responses among the rosette rather than simultaneous activation of individual cells (Figure 3G). Thus, choanoflagellate rosettes reveal how cell-autonomous sensory responses are transformed into coordinated multicellular behavior, establishing them as a valuable model for understanding the evolution of multicellular signal processing.

### TRPW1 has conserved and specialized ion channel domains

Given the sensory function of TRPW1 and its relationship to animal sensory receptors, we used TRPW1 to probe the structural basis of sensory receptor evolution. We first used single-particle cryo-electron microscopy (cryo-EM) to determine the TRPW1 structure at 2.37 Å overall resolution (Figure S4). TRPW1 subunits form a tetrameric complex with a transmembrane domain architecture shared by all members of the voltage-gated superfamily of ion channels, including voltage-gated potassium channels (K_V_) that date to the last universal common ancestor of all cellular life (*23*), and more recently derived TRP channels (Figure 4A-D). Other features of TRPW1 are shared with type I TRP channels, including the “domain-swapped” arrangement around the central ion channel pore (*24*), the signature “TRP helix” lining the cytoplasmic side of the transmembrane region (*25*), and relatively limited extracellular surface, in agreement with its activation by hydrophobic molecules (Figure 4A-D). TRPW also exhibits a C-terminal coiled coil near the channel pore, which is present in many type I TRP families, but absent in type II TRP channels (TRPML and TRPP), which lack many features of sensory type I TRP channels (Figure 4A-D).

**Figure 4:**
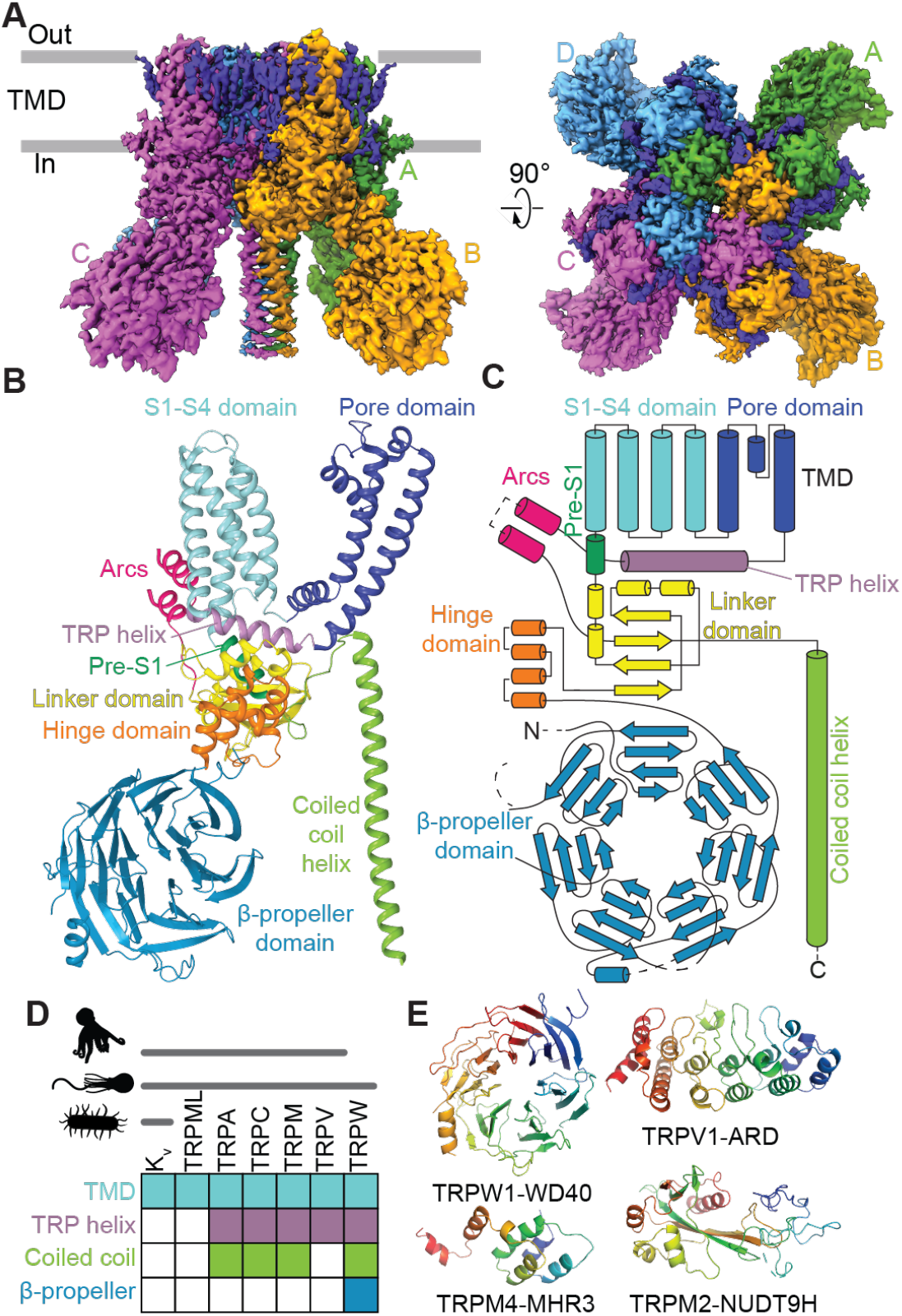
TRPW1 has conserved and specialized ion channel domains. **(A)** Electron density representation of the tetrameric structure of TRPW1 as revealed by cryo-electron microscopy. TRPW1 consists of a transmembrane domain (TMD) and a cytoplasmic domain with WDR β-propellers. The four subunits A-D are colored differently, with membrane lipids additionally shown in dark purple. The top-down view shows the domain-swapped architecture typical of TRP channels and the ion conduction pore in the center of the tetramer. **(B)** Ribbon diagram of a single subunit of TRPW1. The S1-S4 domain and pore domain constitute the transmembrane elements. The Pre-S1, TRP helix and coiled coil helix are common structural elements of type I TRP channels. The seven WD40 domains of TRPW1 fold into a β-propellers architecture. **(C)** Cartoon domain organization of a single subunit of TRPW1, with analogous elements and labeling to (B). **(D)** TRPW1 shows features that are common to many ion channel families across life (e.g. TMD found in K_V_, TRPML), features found in most type I TRP channels (TRP helix, C-terminal coiled coil), and features unique to the TRPW family (WD40 β-propeller). **(E)** Comparison of cytoplasmic domains found in diverse TRP channels, including WDR domains specific to TRPW.

The most distinctive architectural feature of TRPW1 is its N-terminal WD40 repeats. The cryo-EM structure shows that the WD40 repeats of each TRPW1 subunit form an intracellular 7-blade β-propeller WD40 repeat (WDR) domain, with each WD40 repeat contributing one blade (Figure 4A-D). Interestingly, some TRPW family members in *S. rosetta* and other choanoflagellates also contain ankyrin repeat domains between the WD40 repeats and the transmembrane domain (Figure S1E). Ankyrin repeat domains are also found in other type I TRP channels, including TRPA and TRPV. Together with the diversity of N-terminal structures across animal TRP channels, the presence of the WDR suggests that the TRPW modular architecture could facilitate functional diversification and adaptation (Figure 4E, S1E). Indeed, β-propeller domains are known to be modular protein-protein interaction surfaces, including within cilia or flagella (*26–28*), and therefore could link TRPW1 activation to intracellular transduction and flagellar regulation.

### TRPW1 agonist binding reveals ancestral features of sensory receptors

We next asked whether the chemosensory mechanisms of TRPW1 could reveal ancestral features of animal sensory receptors. First, we resolved an additional cryo-EM structure of TRPW1 bound to the agonist RS (Figure 5A,B). Consistent with its activation properties, binding to RS resulted in a widening of the ion conduction pore of TRPW1, suggesting a “pre-activated” state (*29, 30*) (Figure S5A-C). RS was bound in a transmembrane site between S1-S4 and S5-S6 of adjacent subunits, with the TRP helix and S4-S5 linker forming the floor of the pocket (Figure 5A,B). Mutation of residues E828, R898, or Q927 in S1-S4, or R950 in the S4-S5 linker abolished RS-evoked activity, demonstrating the importance of this site for ligand binding and activation (Figure 5C, Figure S5D). Furthermore, substitution of I949W was predicted to enhance the fit of RS and increased apparent affinity by ten-fold (Figure S5E). Importantly, some of these mutations did not preclude activation of TRPW1 by other synthetic agonists, demonstrating that these point mutations did not block general TRPW1 function (Figure S5D). Mutagenesis of R950, R898, and Q927 abolished 15M-9E-evoked activity, supporting a similar binding position as RS (Figure 5C). However, the simple fatty acid structure of 15M-9E precluded experimental structural studies, so we used molecular dynamics simulations to place 15M-9E ligands in this site. We observed that in all replicates, 15M-9E remained stably bound over 500 ns simulations, coordinated by salt bridges with R950 and R1063 and a hydrogen bond with W891, with the fatty tail occupying the hydrophobic cavity of the pocket (Figure 5D, Figure S6A-C). Intriguingly, the binding pocket of TRPW1 is structurally similar to the “vanilloid binding pocket” that the mammalian pain receptor TRPV1 uses to coordinate capsaicin, the “spicy” molecule of *Capsicum* chili peppers (*29*), and lipids involved in inflammatory signaling (*31*–*33*) (Figure 5E). To-gether with their sequence and structural similarities, the shared binding pocket suggests that TRPWs are closely related to sensory receptors in animals. To explore whether TRPWs predate sensory TRP channels in animals, we searched 993 eukaryotic proteomes (*34*) and found the TRPW architecture across diverse clades (Figure 5F). We extracted ion channel domains from candidate orthologs in euglenids, ciliates, diatoms, and chlorophyte algae and inferred phylogenetic trees, finding strong support for their inclusion in the TPRW clade (Figure S7). Furthermore, many of these orthologs also included an ankyrin repeat domain between the WD40 repeats and the ion channel domain, strengthening the evolutionary relationship of these receptors (Figure S7C). Together, these analyses support the emergence of TRPWs at or near the base of the eukaryotic tree (*35*). The type I TRP channels TRPV, TRPA, TRPC and TRPM, have so far mainly been detected in animals and their closest protist relatives (*36, 37*) (Figure 5G). Our phylogenetic analysis did not support the only known reported exceptions: two sequences in dinoflagellates (*38*), one sequence from chlorophytes (*39*), and one sequence from apusozoans (*37*) (Figure S7B). Given the shared features of TRPW with animal sensory TRPs that have not been described in other families like TRPP and TRPML, TRPW is positioned as an ancient ion channel family that can help explain the origin of animal somatosensory receptors.

**Figure 5:**
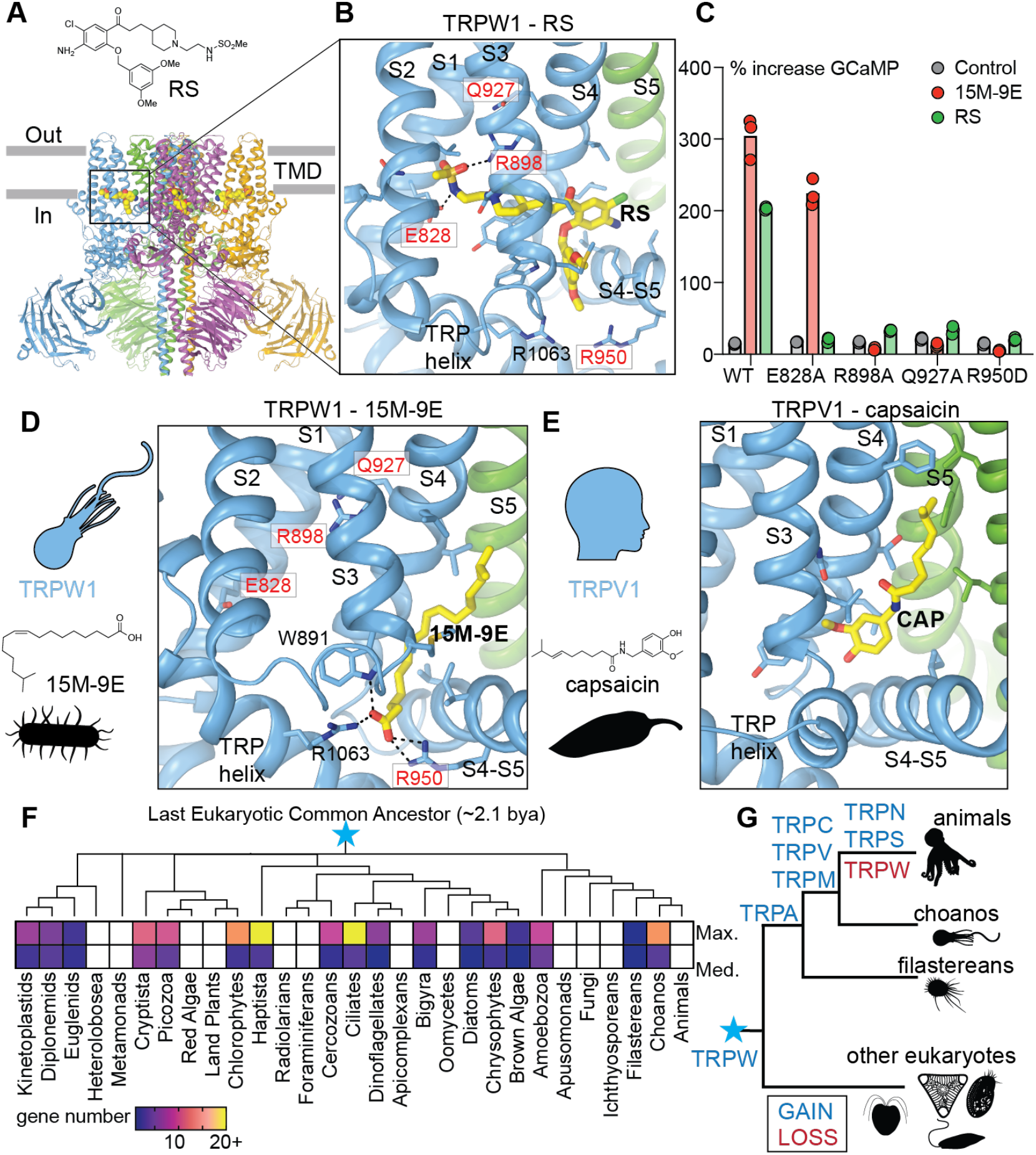
TRPW1 agonist binding reveals ancestral features of sensory TRP channels. **(A)** Cryo-electron microscopy of TRPW1 with the synthetic agonist RS39604 (RS) shows agonist binding near the cytoplasmic leaflet of the plasma membrane at the interface of adjacent subunits. **(B**) Zoom-in of RS binding to TRPW1, positioned between the S1-S4 bundle, TRP helix, and S4-S5 linker of one subunit (colored in blue) and the S5 helix of an adjacent subunit (colored in green). Residues in red were selected for mutagenesis experiments. **(C)** Mutation of E828A is selective for RS activation, while three other mutations – R898A, Q927A, and R950D – abolish activation by both RS and 15M-9E, supporting a similar binding site for these molecules in TRPW1. See Figures S5D-E for further mutagenesis results with an array of TRPW1 agonists. n = 3 Ca^2+^ imaging replicates. **(D)** Molecular modeling suggests that 15M-9E binds near the same pocket as RS. In this pose, the carboxylic acid group of 15M-9E forms salt bridges with R1063 and R950, which are required for 15M-9E-evoked activity. **(E)** The lipid binding pocket in TRPW1 is analogous to the “vanilloid binding pocket” for capsaicin in animal TRPV1. **(F)** TRPW domain architectures are found across eukaryotic groups with maximum and median gene counts per species shown for each clade. Phylogenetic analysis confirms TRPW orthologs across eukaryotic diversity (Figure S7). **(G)** Model of gene family gain and loss indicating that TRPW is ancient but still found in choanoflagellates and other protistan lineages.

## Discussion

TRPW arose more than 2 billion years ago, at or close to the last eukaryotic common ancestor. Eukaryotes emerged in bacteria-rich environments, where early sensory receptors were likely tuned to bacterial cues marking food, danger, competitors, or symbiotic partners. TRPW1 sensitivity to lipids from bacterial prey suggests that bacterial sensation could be one ancestral function of eukaryotic TRP channels. Indeed, the TRPW1 binding pocket resembles the vanilloid binding pocket in TRPV1, which is also a regulatory site in other sensory TRP channels (*29, 40, 41*). Because hydrophobic bacterial lipids diffuse poorly in aquatic environments, they are likely sensed by direct contact with membranes or outer membrane vesicles. A similar logic applies to microbial sensing in octopus chemotactile receptors and TRP channels in the vertebrate gut, suggesting that contact-dependent chemosensation of bacteria has deep evolutionary origins and represents a conserved strategy for extracting information from surfaces (*42–44*).

*S. rosetta* localizes TRPW1 to the flagellum, an ancient eukaryotic organelle that combines environmental sampling with motility, including in animal cells (*16, 45–47*). This arrangement places receptor activation and motor control in the same compartment, allowing for rapid conversion of brief contact with bacterial cues into localized calcium influx, excitation, and immediate behavioral output. Compartmentalization is essential in a unicellular organism, which must transduce distinct environmental cues into appropriate cellular behaviors. Interestingly, the eukaryotic distribution of the TRPW family shows that it has been maintained mostly in free-living protistan lineages, but lost in obligately multicellular clades (animals, land plants) and parasitic lineages (apicomplexans, oomycetes), therefore correlating with a protistan mode of flagellar locomotion. A similar pattern of gene retention in organisms with cilia or flagella was previously observed for voltage-gated calcium channels, which have been reported to regulate flagellar activity in protists (*18, 48*). Supporting this, the flagellum of the unicellular green alga *Chlamydomonas reinhardtii* contains both a voltage-gated calcium channel and a TRP channel, which is required for mechanosensory behaviors (*39, 48, 49*). Our expanded phylogenetic analysis now shows that this TRP channel belongs to the novel TRPW clade, indicating that sensory roles for TRPWs extend beyond choanoflagellates to other protists.

Multicellular choano rosettes extend the logic of compartmentalized sensation within single cells to multicellular co-ordination. TRPW1 agonists trigger synchronized transitions between flagellar beating and arrest across rosette cells, transforming sensation by individual flagella into a coordinated colony-level response (*21*). This organization can be seen as a pre-neuronal solution to the general challenge of coupling localized sensory detection to signal propagation and coordinated behavior across networks of cells. Consistent with this idea, stimulus-evoked coordination in the choanoflagellate *Choanoeca flexa* suggests that multicellular choanoflagellates have evolved diverse strategies for integrating environmental information across colonies (*50*). Because choanoflagellates can be studied as both single cells and multicellular organisms, they provide a unique system for asking how sensory receptors (e.g. TRPW), organellar signaling compartments (e.g. the flagellum), and intercellular communication (e.g. in rosettes) were organized during the emergence of multicellular behavior.

## Acknowledgements

We thank: Douglas Richardson and the Harvard Center for Biological Imaging for consultation, training, and access to microscopes; the Harvard Bauer Core for access to equipment; and Brittany Walsh for help with choano care. This work was supported by an NIH NRSa to MCC and a Human Frontier Science Program (HFSP) Award to JN, IAT, and AIS. Access to computational resources of HPC facilities at NRU HSE University was gratefully appreciated. Nicole King, David Julius, Thibaut Brunet, Rich Losick, Rebecka Sepela, Anastasiia Sukalskaia, Wendy Valencia-Montoya, Alain Garcia de Las Bayonas, and Erika López Alfonzo provided helpful feedback on drafts of the manuscript.

## Author contributions

MCC and NWB conceptualized the study. MCC, JN, YHS, RS, JC, IAT, IIV, YAT, RGE, and AIS contributed to molecular, cellular, structural and behavioral experiments. All authors were involved with writing or reviewing the manuscript.

## Data and materials availability

The cryo-EM density maps for TRPW1-apo (accession codes EMD-77243 for consensus, EMD-77244 for WDR-focused and EMD-77241 for composite maps) and TRPW1-RS39604 (accession codes EMD-77245 for consensus, EMD-77246 for WDR-focused, EMD-77247 for coiled-coil-focused and EMD-77242 for composite maps) have been deposited in the Electron Microscopy Data Bank (EMDB). The atomic coordinates have been deposited in the Protein Data Bank (PDB) with the accession codes 35WH (TRPW1-apo) and 35WI (TRPW1-RS39604).

## References

1. M. Sachkova, V. Modepalli, M. Kittelmann, The deep evolutionary roots of the nervous system. Annu. Rev. Neurosci. 48, 311–329 (2025).

2. K. Y. Wan, Biophysics of protist behaviour. Curr. Biol. 34, R981–R986 (2024).

3. N. Ros-Rocher, T. Brunet, What is it like to be a choanoflagellate? Sensation, processing and behavior in the closest unicellular relatives of animals. Anim. Cogn., doi: 10.1007/s10071-023-01776-z (2023).

4. M. J. Dayel, R. A. Alegado, S. R. Fairclough, T. C. Levin, S. A. Nichols, K. McDonald, N. King, Cell differentiation and morphogenesis in the colony-forming choanoflagellate Salpingoeca rosetta. Dev. Biol. 357, 73–82 (2011).

5. N. Ros-Rocher, J. Reyes-Rivera, U. Horo, C. Combredet, Y. Foroughijabbari, B. T. Larson, M. C. Coyle, E. A. T. Houtepen, M. J. A. Vermeij, J. L. Steenwyk, T. Brunet, Clonal-aggregative multicellularity tuned by salinity in a choanoflagellate. Nature 651, 974–985 (2026).

6. B. S. C. Leadbeater, The Choanoflagellates (Cambridge University Press, 2015).

7. M. J. Dayel, N. King, Prey capture and phagocytosis in the choanoflagellate Salpingoeca rosetta. PLoS One 9, e95577 (2014).

8. R. A. Alegado, L. W. Brown, S. Cao, R. K. Dermenjian, R. Zuzow, S. R. Fairclough, J. Clardy, N. King, A bacterial sulfonolipid triggers multicellular development in the closest living relatives of animals. Elife 1, e00013 (2012).

9. A. Woznica, A. M. Cantley, C. Beemelmanns, E. Freinkman, J. Clardy, N. King, Bacterial lipids activate, synergize, and inhibit a developmental switch in choanoflagellates. Proc. Natl. Acad. Sci. U. S. A. 113, 7894–7899 (2016).

10. W. A. Valencia-Montoya, N. E. Pierce, N. W. Bellono, Evolution of sensory receptors. Annu. Rev. Cell Dev. Biol. 40, 353–379 (2024).

11. M. M. Diver, J. V. Lin King, D. Julius, Y. Cheng, Sensory TRP channels in three dimensions. Annu. Rev. Biochem. 91, 629–649 (2022).

12. R. Juodeikis, S. R. Carding, Outer membrane vesicles: Biogenesis, functions, and issues. Microbiol. Mol. Biol. Rev. 86, e0003222 (2022).

13. H. Götzke, M. Kilisch, M. Martínez-Carranza, S. Sograte-Idrissi, A. Rajavel, T. Schlichthaerle, N. Engels, R. Jungmann, P. Stenmark, F. Opazo, S. Frey, The ALFA-tag is a highly versatile tool for nanobody-based bioscience applications. Nat. Commun. 10, 4403 (2019).

14. M. C. Coyle, A. M. Tajima, F. Leon, S. P. Choksi, A. Yang, S. Espinoza, T. R. Hughes, J. F. Reiter, D. S. Booth, N. King, An RFX transcription factor regulates ciliogenesis in the closest living relatives of animals. Curr. Biol., doi: 10.1016/j.cub.2023.07.022 (2023).

15. D. R. Mitchell, The evolution of eukaryotic cilia and flagella as motile and sensory organelles. Adv. Exp. Med. Biol. 607, 130–140 (2007).

16. R. A. Bloodgood, Sensory reception is an attribute of both primary cilia and motile cilia. J. Cell Sci. 123, 505–509 (2010).

17. Z. Carvalho-Santos, J. Azimzadeh, J. B. Pereira-Leal, M. Bettencourt-Dias, Evolution: Tracing the origins of centrioles, cilia, and flagella. J. Cell Biol. 194, 165–175 (2011).

18. T. Brunet, D. Arendt, From damage response to action potentials: early evolution of neural and contractile modules in stem eukaryotes. Philos. Trans. R. Soc. Lond. B Biol. Sci. 371, 20150043 (2016).

19. K. Mizuno, K. Shiba, M. Okai, Y. Takahashi, Y. Shitaka, K. Oiwa, M. Tanokura, K. Inaba, Calaxin drives sperm chemotaxis by Ca2+-mediated direct modulation of a dynein motor. Proc. Natl. Acad. Sci. U. S. A. 109, 20497–20502 (2012).

20. D. Laundon, B. T. Larson, K. McDonald, N. King, P. Burkhardt, The architecture of cell differentiation in choanoflagellates and sponge choanocytes. PLoS Biol. 17, e3000226 (2019).

21. J. Colgren, P. Burkhardt, Electrical signaling and coordinated behavior in the closest relative of animals. Sci. Adv. 11, eadr7434 (2025).

22. E. A. Lumpkin, M. J. Caterina, Mechanisms of sensory transduction in the skin. Nature 445, 858–865 (2007).

23. F. H. Yu, W. A. Catterall, The VGL-chanome: a protein superfamily specialized for electrical signaling and ionic homeostasis. Sci. STKE 2004, re15 (2004).

24. N. J. Himmel, D. N. Cox, Transient receptor potential channels: current perspectives on evolution, structure, function and nomenclature. Proc. Biol. Sci. 287, 20201309 (2020).

25. D. Granata, V. Carnevale, J. C. Opazo, S. E. Brauchi, Sequence and structural conservation reveal fingerprint residues in TRP channels. Elife (2022).

26. C. U. Stirnimann, E. Petsalaki, R. B. Russell, C. W. Müller, WD40 proteins propel cellular networks. Trends Biochem. Sci. 35, 565–574 (2010).

27. A. Accogli, S. Shakya, T. Yang, C. Insinna, S. Y. Kim, D. Bell, K. R. Butov, M. Severino, M. Niceta, M. Scala, H. S. Lee, T. Yoo, J. Stauffer, H. Zhao, C. Fiorillo, M. Pedemonte, M. C. Diana, S. Baldassari, V. Zakharova, A. Shcherbina, Y. Rodina, C. Fagerberg, L. S. Roos, J. Wierzba, A. Dobosz, A. Gerard, L. Potocki, J. A. Rosenfeld, S. R. Lalani, T. M. Scott, D. Scott, M. S. Azamian, R. Louie, H. W. Moore, N. L. Champaigne, G. Hollingsworth, A. Torella, V. Nigro, R. Ploski, V. Salpietro, F. Zara, S. Pizzi, G. Chillemi, M. Ognibene, E. Cooney, J. Do, A. Linnemann, M. J. Larsen, S. Specht, K. J. Walters, H.-J. Choi, M. Choi, M. Tartaglia, P. Youkharibache, J.-H. Chae, V. Capra, S.-G. Park, C. J. Westlake, Variants in the WDR44 WD40-repeat domain cause a spectrum of ciliopathy by impairing ciliogenesis initiation. Nat. Commun. 15, 365 (2024).

28. Y. Kim, S. H. Kim, WD40-repeat proteins in ciliopathies and congenital disorders of endocrine system. Endocrinology and Metabolism 35, 494–506 (2020).

29. E. Cao, M. Liao, Y. Cheng, D. Julius, TRPV1 structures in distinct conformations reveal activation mechanisms. Nature 504, 113–118 (2013).

30. Y. Yin, S. C. Le, A. L. Hsu, M. J. Borgnia, H. Yang, S.-Y. Lee, Structural basis of cooling agent and lipid sensing by the cold-activated TRPM8 channel. Science 363, eaav9334 (2019).

31. Y. Gao, E. Cao, D. Julius, Y. Cheng, TRPV1 structures in nanodiscs reveal mechanisms of ligand and lipid action. Nature 534, 347–351 (2016).

32. W. R. Arnold, A. Mancino, F. R. Moss 3rd, A. Frost, D. Julius, Y. Cheng, Structural basis of TRPV1 modulation by endogenous bioactive lipids. Nat. Struct. Mol. Biol. 31, 1377–1385 (2024).

33. Y. A. Trofimov, N. A. Krylov, A. S. Minakov, K. D. Nadezhdin, A. Neuberger, A. I. Sobolevsky, R. G. Efremov, Dynamic molecular portraits of ion-conducting pores characterize functional states of TRPV channels. Commun. Chem. 7, 119 (2024).

34. D. J. Richter, C. Berney, J. F. H. Strassert, Y.-P. Poh, E. K. Herman, S. A. Muñoz-Gómez, J. G. Wideman, F. Burki, C. de Vargas, EukProt: a database of genome-scale predicted proteins across the diversity of eukaryotes. Peer Community Journal, doi: 10.24072/pcjournal.173 (2022).

35. G. Torruella, L. J. Galindo, D. Moreira, P. López-García, Phylogenomics of neglected flagellated protists supports a revised eukaryotic tree of life. Curr. Biol. 35, 198–207.e4 (2025).

36. G. Peng, X. Shi, T. Kadowaki, Evolution of TRP channels inferred by their classification in diverse animal species. Mol. Phylogenet. Evol. 84, 145–157 (2015).

37. X. Cai, D. E. Clapham, Ancestral Ca2+ signaling machinery in early animal and fungal evolution. Mol. Biol. Evol. 29, 91–100 (2012).

38. J. B. Lindström, N. T. Pierce, M. I. Latz, Role of TRP channels in dinoflagellate mechanotransduction. Biol. Bull. 233, 151–167 (2017).

39. K. Fujiu, Y. Nakayama, H. Iida, M. Sokabe, K. Yoshimura, Mechanoreception in motile flagella of Chlamydomonas. Nat. Cell Biol. 13, 630–632 (2011).

40. C. Liu, R. Reese, S. Vu, L. Rougé, S. D. Shields, S. Kakiuchi-Kiyota, H. Chen, K. Johnson, Y. P. Shi, T. Chernov-Rogan, D. M. Z. Greiner, P. B. Kohli, D. Hackos, B. Brillantes, C. Tam, T. Li, J. Wang, B. Safina, S. Magnuson, M. Volgraf, J. Payandeh, J. Zheng, A. Rohou, J. Chen, A non-covalent ligand reveals biased agonism of the TRPA1 ion channel. Neuron 109, 273–284.e4 (2021).

41. Q. Tang, W. Guo, L. Zheng, J.-X. Wu, M. Liu, X. Zhou, X. Zhang, L. Chen, Structure of the receptor-activated human TRPC6 and TRPC3 ion channels. Cell Res. 28, 746–755 (2018).

42. R. J. Sepela, H. Jiang, Y.-H. Shin, T. L. Hautala, J. Clardy, R. E. Hibbs, N. W. Bellono, Environmental microbiomes drive chemotactile sensation in octopus. Cell 188, 4849–4860.e21 (2025).

43. N. W. Bellono, J. R. Bayrer, D. B. Leitch, J. Castro, C. Zhang, T. A. O’Donnell, S. M. Brierley, H. A. Ingraham, D. Julius, Enterochromaffin cells are gut chemosensors that couple to sensory neural pathways. Cell 170, 185–198.e16 (2017).

44. L. Ye, M. Bae, C. D. Cassilly, S. V. Jabba, D. W. Thorpe, A. M. Martin, H.-Y. Lu, J. Wang, J. D. Thompson, C. R. Lickwar, K. D. Poss, D. J. Keating, S.-E. Jordt, J. Clardy, R. A. Liddle, J. F. Rawls, Enteroendocrine cells sense bacterial tryptophan catabolites to activate enteric and vagal neuronal pathways. Cell Host Microbe 29, 179–196.e9 (2021).

45. G. Jékely, D. Arendt, Evolution of intraflagellar transport from coated vesicles and autogenous origin of the eukaryotic cilium. Bioessays 28, 191–198 (2006).

46. M. Delling, P. G. DeCaen, J. F. Doerner, S. Febvay, D. E. Clapham, Primary cilia are specialized calcium signalling organelles. Nature 504, 311–314 (2013).

47. K. I. Hilgendorf, B. R. Myers, J. F. Reiter, Emerging mechanistic understanding of cilia function in cellular signalling. Nat. Rev. Mol. Cell Biol. 25, 555–573 (2024).

48. K. Fujiu, Y. Nakayama, A. Yanagisawa, M. Sokabe, K. Yoshimura, Chlamydomonas CAV2 encodes a voltage-dependent calcium channel required for the flagellar waveform conversion. Curr. Biol. 19, 133–139 (2009).

49. D. Oshima, M. Yoshida, K. Saga, N. Ito, M. Tsuji, A. Isu, N. Watanabe, K.-I. Wakabayashi, K. Yoshimura, Mechanoresponses mediated by the TRP11 channel in cilia of Chlamydomonas reinhardtii. iScience 26, 107926 (2023).

50. T. Brunet, B. T. Larson, T. A. Linden, M. J. A. Vermeij, K. McDonald, N. King, Light-regulated collective contractility in a multicellular choanoflagellate. Science 366, 326–334 (2019).

