## Supplementary Info for "An ancient receptor family illuminates the evolution of animal sensation"

#### The PDF file includes:

Materials and Methods

Supplementary Figures 1-7

Supplementary Movies 1-3

Supplementary Data Files 1-6

Supplementary Tables 1-4

References

#### Materials and Methods

##### Bioinformatics

Ortholog identification

To identify gene family expansions in choanoflagellates relative to their sister lineage (animals) as well as outgroups (filastereans, ichthyosporeans), diverse proteomes were downloaded from the EukProt database (*1*). The dataset included:

10 choanoflagellates:

EP00035_Acanthoeca_spectabilis, EP00037_Helgoeca_nana, EP00039_Didymoeca_costata, EP00040_Diaphanoeca_grandis, EP00041_Stephanoeca_diplocostata, EP00042_Codosiga_hollandica, EP00044_Salpingoeca_rosetta, EP00046_Monosiga_brevicollis, EP00050_Salpingoeca_kvevrii, EP00054_Salpingoeca_dolichothecata

10 animals:

EP00057_Strongylocentrotus_purpuratus, EP00067_Danio_rerio, EP00090_Calanus_glacialis, EP00106_Octopus_bimaculoides, EP00110_Nematostella_vectensis, EP00111_Clytia_hemisphaerica, EP00114_Trichoplax_sp_H2, EP00115_Mnemiopsis_leidyi, EP00118_Oscarella_pearsei, EP00119_Amphimedon_queenslandica

2 filastereans:

EP00120_Capsaspora_owczarzaki, EP01137_Pigoraptor_chileana

2 ichthyosporeans:

EP00124_Ichthyophonus_hoferi, EP00125_Sphaeroforma_arctica

Orthogroups were assigned using OrthoFinder v2.5.5 (*2*), using the -og flag to stop after orthogroup inference. Out of 708,653 genes, 78.4% were assigned among 57,468 orthogroups. This dataset was manually inspected for gene families with higher copy number in choanoflagellates than in animals or outgroups, annotating candidate gene families for the presence of transmembrane regions and other protein domains. This approach identified OG_594, an orthogroup containing many genes that featured an ion channel domain (Pfam profile PF00520) and WD40 repeats (Pfam profile PF00400) (*3*). This orthogroup contained 16 members from *Salpingoeca roset*ta, all with an ion channel transmembrane domain, and 13 of which additionally showed one or more WD40 repeats.

Phylogenetic tree of PF00520 genes from *S. rosetta*

To place the novel family in phylogenetic context, the PF00520 Pfam domain, representing the ion channel found in TRP channels, voltage-gated channels, and other channels (*4*), was queried against the *S. rosetta* proteome using hmmsearch from the HMMer toolkit using the default threshold (*5*). Hits were manually annotated for gene family affiliation using BLAST (*6*) and protein domain architecture. This revealed one more gene, PTSG_06082, containing an ion channel domain and WD40 repeats, which was not found in the original orthogroup OG_594.

The ion channel sequences for each hit were extracted (for genes with multiple 6TM domains, particularly two-pore channels and $Na_V$/$Ca_V$, the first 6TM domain was selected), aligned with MAFFT v7.526 (*7*), trimmed with ClipKIT v1.3.0 (*8*), and a maximum-likelihood phylogenetic tree was constructed using IQTREE v2.4.0 (*9*, *10*). The resulting tree placed all 16 members of OG_594 in the same monophyletic group with high statistical support (91/94 for UFboot/SH-aLRT). This suggested that the three genes in OG_594 without WD40 domains are likely evolutionarily related to the rest of the group. The phylogeny placed PTSG_06082, the WD40-containing ion channel not originally identified in OG_594, in a sister clade to the rest of the family, along with PTSG_00805, which has been identified (and further supported by analysis here) as a likely TRPV ortholog (*11*, *12*). Based on the presence of WD40 repeats and ion channel transmembrane domains, and the near phylogenetic affiliation with the rest of the TRPW group, PTSG_06082 was included as a TRPW sequence. Finally, the phylogenetic analysis showed one gene, PTSG_11274 in the TRPW clade, which lacks WD40 repeats and was not identified in OG_594. Furthermore, this gene contains a predicted SLOG domain and Nudt9 domain, typical of TRPM family members (*13*) and found in many *S. rosetta* TRPM homologs that grouped as a separate clade in our phylogeny. Based on its domain architecture, PTSG_11274 was not included as a TRPW, but was instead assigned as a divergent TRPM ortholog based on its domain architecture. This left a final set of 17 TRPW orthologs, 14 of which contain WD40 repeats, and 2 of which additionally encode ankyrin repeats, which are common in other TRP channel classes such as TRPV, TRPA, and TRPC channels. Altogether, the TRPWs grouped together with *S. rosetta* TRPMs, TRPAs, TRPV and TRPC in a clade with 82/92 support by UFboot/SH-aLRT. This clade included one other PF005200-containing sequence, PTSG_07737, which contains a cyclic nucleotide binding domain and was placed as a particularly long branch in the phylogeny. This gene is more likely evolutionarily related to the CNG family, a different sub-family of the voltage-gated superfamily (*4*). For display in Figure 1B, all branches with <70% support by UFboot are collapsed into palintomies.

<u>Search for WD40+ion channel domain architecture across eukaryotes</u>

The unique and diagnostic architecture of TRPW sequences (WD40 repeats + ion channel transmembrane domain) led us to search for potential orthologs more broadly across eukaryotic diversity. Pfam profiles for the ion channel transmembrane domain (PF00520) and the WD40 repeats (PF00400) were queried against the entire EukProt proteome collection, consisting of 993 proteomes curated to span the taxonomic diversity of eukaryotes. Default threshold criteria were used for calling hits, then proteins were identified that contained hits to both domains. Proteins with the WD40 + ion channel architecture were found in 184 distinct taxa, occurring repeatedly in closely related taxa of different groups. Most taxa with putative orthologs are protists, as no hits were found in any animals, land plants, fungi, or red algae, although some brown algae contained putative orthologs (e.g. *Ectocarpus siliculosus*). Groups with a particular rich diversity of TRPW orthologs included choanoflagellates, ciliates, chlorophyte algae, and haptists. For display (Figure 1C, Figure 5F), the maximum and median number of genes with this domain architecture were shown for each group, considering those taxa for which hits were found (e.g. the median for ciliates is the median number of TRPW genes in all TRPW-containing ciliate species, not in all ciliates queried).

Two false positive hits were initially identified in animals and closely investigated. (1) The ctenophore sequence EP00116_Pleurobrachia_bachei_P001090 contains WD40 repeats in the C-terminal portion of the protein, which altogether shows high percent identity (>65%) to dynein intermediate chain 3 across diverse animals. This protein additionally contains a BTB/POZ domain, common in voltage-gated potassium channels that also have the PF00520 domain. Therefore, this protein is most likely a fusion (real or due to genome assembly artifacts) between a voltage-gated potassium channel and a dynein intermediate chain and does not resemble the domain architecture of choanoflagellate TRPWs. (2) The nematode sequence EP00080_Pristionchus_pacificus_P007218 contains two ion channel domains and has accordingly been annotated as a homolog of *C. elegans* twk-25, a two-pore domain potassium channel (K2P). The N-terminal WD40 repeats of this protein may be due to a fusion (again, real or due to genome assembly artifacts) of a WD40 repeat-containing protein and a K2P channel. In contrast to these proteins, domain architecture hits from diverse protists (ciliates, filastereans, diatoms, chlorophytes) resembled choanoflagellate TRPWs (Figure S7C). Together, these data indicate a secondary loss of TRPW in the animal lineage.

<u>Phylogenetic tree of animal and choanoflagellate TRP channel sequences</u>

To more thoroughly understand the distribution and phylogenetic relationships of TRP channels within choanoflagellates, orthogroups containing previously annotated *S. rosetta* TRP channels (*11*, *12*), including TRPA, TRPC, TRPV, and TRPM channels were identified. Choano representatives of these four orthogroups were combined with the orthogroup for candidate TRPWs in a single FASTA file. These sequences were queried with PF00520 using hmmsearch, and ion channel domains were extracted, aligned with MAFFT and trimmed with ClipKIT, followed by tree inference with IQTREE. The resulting tree nicely separated five families of TRP channels, with accessory domains supporting channel affiliation (ankyrin for TRPA, TRPV, TRPC, WD40 for TRPW, TRPM2, Nudix and SLOG domains for TRPM). For each of these phylogenetically validated families, a custom HMM profile was built using hmmbuild after extracting ion channel sequences and alignment with MAFFT.

Next, all available choanoflagellate proteomes on EukProt with BUSCO scores >60% were queried with hmmsearch using the custom profiles for the individual TRP channels. These were filtered for only the highest-confidence hits, using domain scores >30, predicted protein lengths >100 amino acids, and only including predicted full-length proteins (with in-frame start and stop codons). A further tree was inferred for these high-confidence hits (using the ion channel domain) across all available choano data. The resulting tree identified high-confidence groups, with previous *S. rosetta* orthologs for TRPA, TRPC, TRPM and TRPV falling into distinct groups with >95% bootstrap support and consistent domain architectures. TRPW was well-supported as a distinct group with high statistical support (97% UF-boot, 97 SH-aLRT). Two small clades outside of these main groups appeared with predicted PKD/polycystin domains. These were removed from further analysis to narrow our focus on type I TRP channels, but these may represent TRPP and TRPML orthologs.

To ascertain the phylogenetic affinity of choanoflagellate TRP channel families with animal TRP channels, a taxonomically diverse set of animal proteomes was selected from EukProt:
EP00057_Strongylocentrotus_purpuratus, EP00058_Branchiostoma_floridae,
EP00059_Ciona_intestinalis, EP00063_Salmo_salar, EP00074_Homo_sapiens,
EP00081_Caenorhabditis_elegans, EP00098_Tribolium_castaneum,
EP00099_Drosophila_melanogaster, EP00103_Capitella_teleta,
EP00106_Octopus_bimaculoides, EP00107_Lottia_gigantea, EP00110_Nematostella_vectensis,
EP00111_Clytia_hemisphaerica, EP00113_Trichoplax_adhaerens,
EP00115_Mnemiopsis_leidyi, EP00118_Oscarella_pearsei,
EP00119_Amphimedon_queenslandica.

Against these proteomes were queried ion channel domains from the following genes representative of previously recognized animal TRP channel families: human TRPV1, human TRPA1, human TRPM8, human TRPC1, fruit fly nompC, and sea anemone TRPVL (Nemve1|21409|gw.103.50.1). FASTA results for each query were downloaded, concatenated, and CD-HIT was used with a threshold of 0.9 to reduce close isoforms/paralogs. From this collected list of genes, ion channel transmembrane domains were extracted using hmmsearch with PF00520, filtering for the best scoring domain per query sequence and a score >= 21 and length >= 100. CD-HIT was used once more with a threshold of 0.9. For the highly divergent TRPS channels (*13*), the transmembrane domains for 4 gene family members from mollusk, nematode, and lancelet were manually added to the full set. The resulting set of animal TRP channel sequences was combined with the previously assembled choanoflagellate set, aligned with MAFFT, trimmed with ClipKIT, and a maximum-likelihood phylogeny inferred with IQTREE. The resulting tree showed high-confidence clades for choanozoan TRPM/S (96/99 by UFboot-sh-aLRT), choanozoan TRPA (100/100), choanozoan TRPC/N (99/80), choanozoan TRPVL (95/99), and choanoflagellate TRPW (96/95). For TRPV, a high-confidence clade of choanozoan sequences (97/100) was found but included only loricate choano sequences. However, a group of craspedid sequences was found sister to this group with 82/99 support for the combination of these with the loricate+animal TRPVs. These craspedid sequences include a C-terminal domain annotated by Pfam as isochorismatase-like and with highest BLAST similarity to nicotinamidase sequences. These likely represent a divergent ortholog of TRPV in craspedid choanoflagellates which included the fusion of a nicotinamidase domain. For display, all branches with less than 70% support by UFboot were collapsed into palintomies (Figure S1D). Four long-branch sequences with no closely related sequences were trimmed from the final tree: *Acanthoeca spectabilis* P006044, *Bicosta minor* P010219, *Didymoeca costata* P020577, *Clytia hemisphaerica* P015121.

Phylogenetic trees of animal, choanoflagellate, and other eukaryotic TRP channel sequences

First, a subset of the sequences used for the animal+choanoflagellate tree were selected. The list of taxa included was trimmed by removing sequences from *Salmo salar*, *Clytia hemisphaerica*, *Amphimedon queenslandica*, *Trichoplax adhaerens*, *Bicosta minor*, *Microstomoeca roanoka*, *Acanethoecidae* sp. 10tr, *Salpingoeca dolichothecata*, and *Codosiga hollandica*. Also trimmed were any sequences shorter than 180 amino acids and *Didymoeca costata* P020577, which placed inconsistently in previous choanoflagellate gene trees.

Next, sequences from diverse eukaryotic taxa with proteins containing the PF00520 (voltage-gated superfamily ion channel) + PF00400 (WD40) protein domains were added to the choano + animal set. These consisted of sequences from two diatoms (*Chaetoceros debilis*, *Seminavis robusta*), two ciliates (*Fabrea salina, Stylonychia lemnae*), three chlorophyte algae (*Chlamydomonas reinhardtii*, *Volvox carteri*, *Gonium pectorale*), and one euglenid (*Euglena longa*), for a total of 31 additional sequences. HMMer was used to extract the portion of each protein with the ion channel domain. As before, a maximum-likelihood phylogeny was inferred by IQTREE after sequence alignment by MAFFT and trimming by ClipKIT. The resulting tree included all new candidate ion channel+WD40 proteins in a high confidence clade with choanoflagellate TRPWs, with 96/91 support by UFboot/SH-aLRT. Monophyletic groups were inferred for chlorophyte TRPW, diatom TRPW, and ciliate TRPW, with the euglenid sequences branching within the choano TRPW clade. High confidence clades were also inferred from choanozoan TRPC+N, choanozoan TRPM+S, choanozoan TRPA, choanozoan TRPV, and choanozoan TRPVL.

Next, literature was searched for previous reports of protist sequences reported as related to type I TRP channels, finding sequences from chlorophyte algae (*14*, *15*) (reported as TRPV-like), one dinoflagellate (*16*) (reported as TRPM-like) and one apusozoan (*11*) (reported as TRPV-like). The ion channel domains (PF00520 hits) were extracted and added to the previously collected sequence, with alignment and tree inference performed as previously described. Many of the chlorophyte sequences grouped with chlorophyte TRPWs in the TRPW clade, while the rest of the chlorophyte sequences formed a separate clade with no clear relationship to any other sequences. Of the chlorophyte sequences grouping with TRPWs was *C. reinhardtii* TRP11, which lacks WD40 repeats, but was previously reported to localize to the base of the algal flagellum, where it is required for mechanosensitive behaviors (*14*, *17*).

The dinoflagellate and apusozoan sequences showed no clear affiliation, to other sequences in the group or to each other. After collapsing branches with less than 70% UFboot support into palintomies, these other sequences radiated individually from a middle position within the tree (Figure S7B).

**Heterologous expression and physiology**

HEK cell culture and transfection

HEK293T cells (ATCC) were grown in DMEM, 10% fetal bovine serum (PEAK Serum) and 50 IU/mL penicillin and 50 µg/mL streptomycin (Gibco) at 37 °C and 5% $CO_2$. Cells were transfected in Opti-MEM Reduced Serum Media (Gibco) with Lipofectamine 2000 (Invitrogen) for 4-8 hours at 37 °C following the manufacturer's protocol. 1 µg of TRPW1 plasmid was used per 1 mL of transfection mix. For electrophysiology, 0.3 µg of GFP plasmid was added to 1 mL transfection mix with1 µg TRPW1 and incubated in a 35 mm cell culture dish. After transfection, cells were lifted with trypsin and passaged into a fresh 35 mm cell culture dish with 12 mm glass coverslips and standard media. For GCaMP experiments in a 96-well plate, 0.5 µg of GCaMP6s plasmid was added per 1 mL of transfection mix, and 38 µL of transfection mix were added per well of a 96-well plate pre-coated with 0.05 mg/mL poly-D-lysine (Gibco) and washed once with D-PBS. For both electrophysiology and calcium imaging experiments, cells were cultured at 37 °C in fresh media following transfection for 24-48 hours.

Cloning and mutagenesis of TRPW sequences

The coding DNA sequence for TRPW1 and other *S. rosetta* TRPWs was taken from the genome annotation available on Ensembl Protists. Coding sequences were cloned into the pUNIV vector by GenScript, which positions the coding sequence between a T7 promoter and CMV enhancer and *Xenopus* globin 3'UTR. An HA (YPYDVPDYA) tag was added to the C-terminus of the CDS. A separate plasmid was cloned with an additional N-terminal hRho tag (MNGTEGPNFYVPFSNATGVV) to help with surface expression. Once agonists of TRPW1 were identified, responses appeared indistinguishable in cells transfected with either TRPW1-HA or hRho-TRPW1-HA, therefore these constructs were used interchangeably for plate-reader based calcium imaging and electrophysiology experiments. Mutagenesis was done by GenScript, using pUNIV-TRPW1-HA as the base vector. For preparation of further plasmid stocks, plasmids were transformed into XL10-Gold competent cells (Agilent Technologies Cat. No. 200314), grown in liquid LB culture, and DNA was extracted using a Midiprep kit from Zymo (ZymoPURE II Midiprep, Cat. No. D4201).

For structural experiments, cDNA for full-length wild-type *TRPW1* was introduced into *pEG-BacMam* vector for protein expression in mammalian cells (*18*), with its C-terminal region followed by the coding regions for a glycine-threonine-glycine (GTG) linker and the streptavidin affinity tag (the corresponding protein residues WSHPQFEK).

Patch clamp electrophysiology

Whole-cell recordings were conducted from cultured HEK cells in a perfusion chamber (Warner Instruments) using a MultiClamp 700B amplifier (Axon Instruments). The signal was low pass filtered at 1 kHz and acquired at 10 kHz using the Digidata 1550B interface (Axon Instruments). The amplifier was controlled using pCLAMP software (Axon Instruments). Cells were visualized using an inverted fluorescence microscope with DIC contrast (Olympus IX73). Internal solution had the following composition (in mM): 150 $CsMeSO_4$, 5 NaCl, 1 $MgCl_2$, 10 HEPES, 10 EGTA and 10 sucrose. The pH of the internal solution was adjusted to 7.2 with CsOH. Patch pipettes were pulled using a horizontal puller (P-97, Sutter Instrument) from thick-wall borosilicate glass capillaries (Sutter Instrument) to a resistance of 2-5 MOhm. The extracellular solution was mammalian Ringer's, with the following composition (in mM): 140 NaCl, 5 KCl, 10 HEPES, 10 glucose, 2 $CaCl_2$, 2 $MgCl_2$, and adjusted to pH 7.4 with NaOH. Perfusion of chemicals was performed using the SmartSquirt Micro-Perfusion system (Automate Scientific). The voltage-step protocol held a given voltage for 1 second, moving from -120 mV to +120 mV in 20 mV increments, returning to 0 mV potential before and after each step. The voltage-ramp protocol used was a 500 ms ramp from -120 mV to +80 mV. For plotting kinetic data, currents were measured as the average current elicited during a 20 ms window, 40 ms after the start of the 500 ms ramp from −120 to +80 mV. For ion substitution experiments, relative permeability was determined after the substitution of equimolar cations by measuring the shift in $E_{rev}$ from I-V relationships using the above-described voltage-ramp protocol. The extracellular solution contained (in mM): 150 $Na^+$, $Cs^+$, $NMDG^+$, or 100 $Ca^{2+}$ and intracellular solution contained 150 CsCl and 1 CsEGTA, with all solutions buffered with 10 mM HEPES. Permeability ratios were estimated using the Goldman-Hodgkin-Katz (GHK) equation: $P_{Na}/P_{Cs} = (\gamma_{Cs}[Cs^+]_{Cytoplasmic}/\gamma_{Na}[Na^+]_{Luminal})(\exp(E_{rev,Na}-E_{rev,Cs})F/RT))$; $P_{NMDG}/P_{Cs} = (\gamma_{Cs}[Cs^+]_{Cytoplasmic}/\gamma_{Na}[NMDG^+]_{Luminal})(\exp(E_{rev,NMDG}-E_{rev,Cs})F/RT))$;$P_{Ca}/P_{Cs} = (\gamma_{Cs}\,[Cs^+]_{Cytoplasmic}/4\gamma_{Cs}[Ca^{2+}]_{Luminal})(\exp(E_{rev,Ca}-E_{rev,Cs})F/RT))$. $\gamma_{Na}$=0.75; $\gamma_{Cs}$=0.71; $\gamma_{Ca}$=0.29; $\gamma_{NMDG}$=0.80. The agonist for ion substitution experiments was 5 μM 11-deoxy-16-16-dimehtyl prostaglandin E2. Data were analyzed in Igor64 software.

Plate-reader calcium assay

Cells were cultured in clear 96-well cell-culture treated plates and co-transfected with TRPW plasmids and GCaMP6s as described above, with cells between 50 and 80% confluent in the plate. After transfection, fresh media was restored to the cells and they were cultured at 37 °C for 24 hours, at which point the cells were fully confluent. The culture media was replaced by 50 μL per well of mammalian Ringer's solution (see above) and cells kept at room temperature for at least 1h (and no more than 3 h) for signal to reach low baseline levels. Fluorescence measurements on GCaMP expressing cells were performed using the BioTek Synergy Neo2 plate reader (Agilent) using total induced fluorescence emitted from the top. Chemicals were prepared at 3x dilutions in a v-bottom 96-cell plate and 25 μL from this plate were added to the cells after 3 min of baseline measurements using a multi-channel pipettor. Fluorescence in each well was measured every 30 secs for 17 min after drug application and responses quantified as the peak of the calcium trace after chemical application normalized to baseline fluorescence (and reported as % increase), excluding the first 7 time points after drug application when addition of DMSO evokes transient increases in GCaMP activity even in control cells. For the 15M-9E dose-response curve in Figure 2D, low solubility of this ligand required 5.34% DMSO to reach 1 mM drug. This concentration of DMSO causes a higher-amplitude transient spike in GCaMP fluorescence that takes longer to diminish. However, this response was clearly distinguishable from TRPW1-evoked activity which showed sustained GCaMP activity throughout the time-course. Therefore, for this dose-response curve, activity values were reported as % increase of the final timepoint relative to baseline, to distinguish between sustained activity vs. transient activity from the high concentration of DMSO.

For dose-response curves, half-log dilutions were made starting at 300 μM. Curves were fit in Prism 10 software using a four-parameter logistic curve with the model Y=Bottom + (Top-Bottom)/(1+10^((LogEC50-X)*HillSlope).

**Bacterial culture, fractionation, and small molecule discovery**

Preparation of outer membrane vesicles (OMVs) from *Algoriphagus machipongonensis*.

*Algoriphagus machipongonensis* was inoculated into 500 mL of PYG medium (5 g of peptone, 3 g of yeast extract, and 3 mL of glycerol in 1 L of sterilized artificial seawater) and grown at 30 °C for 72 hours in a shaking incubator. Conditioned media was collected by pelleting cells in 50 mL tubes at 3220 x g for 30 mins, followed by decanting the supernatant and filtering twice through 0.22 μm vacuum filters (Corning). The conditioned media was ultra-centrifuged in an Optima L-90K ultracentrifuge (Beckman Coulter) using an SW 32 Ti rotor with 6 tubes of 35 mL conditioned media per tube. These were spun at 37,000 RPM for 3 hours at 4 °C. Supernatant was decanted and each pellet was resuspended in 200 μL of mammalian Ringer's solution and stored at 4 °C.

Bacterial culture for screening

A total of 14 bacterial strains (Table S1) including *Algoriphagus machipongonensis* were initially inoculated into 5 mL of PYG media and cultured for 2 days at 30 °C. The cultures were transferred to 15 mL conical centrifuge tubes and separated into cell pellets and supernatant by centrifugation (4500 rpm at 4 °C for 30 min). To extract intracellular metabolites, the resulting cell pellets were transferred into 4 mL glass vials and disrupted with 2 mL of chloroform-methanol mixture (1:1, v/v) using glass beads and magnetic bars. The mixtures were then filtered through cotton-plugged Pasteur pipettes to remove cell debris, and the filtrates were concentrated under reduced pressure to yield the crude extracts.

Large-scale bacterial culture and extractions

*Algoriphagus machipongonensis* was inoculated into 5 mL of PYG medium and incubated at 30°C for 2 days on a rotary shaker. The entire culture was then used to inoculate 1 L of PYG medium in Ultra Yield® 2.5 L flask for scale-up and incubated under the same conditions. A total of 76 L of liquid culture of *A. machipongonensis* was prepared, and after 4 days of cultivation, the cultures were separated into cell pellets and supernatants by centrifugation (7000 rpm at 4°C for 30 min). The cell pellets were transferred into a 1 L glass bottle and extracted with a 1:1 (v/v) mixture of chloroform-methanol and glass beads by stirring at room temperature overnight. The solvent extract was then filtered through filter paper and dried under low pressure, yielding a crude cell extract.

Activity-guided fractionation and purification of 15M-9E

The crude cell extract was adsorbed onto silica resin and loaded onto a silica column for normal-phase fractionation. Five different solvent systems [chloroform, ethyl acetate-methanol (9:1, v/v), acetone-methanol (9:1, v/v), dichloromethane-methanol (9:1, v/v), and methanol] were used for the fractionation. Fractions were assayed at 100 µg/mL against HEK293Tcells expressing TRPW1 and GCaMP6s, using a plate reader fluorescence assay. This same assay was used at each successive purification step, always with 100 µg/mL of each fraction. The most bioactive fraction on TRPW1 expressing cells was eluted with the ethyl acetate-methanol mixture, was then dried, reconstituted in methanol, and fractionated into 23 fractions by size-exclusion chromatography (Sephadex LH-20, methanol). One of the bioactive Sephadex fractions (fraction 12) was then injected onto a semi-preparative HPLC column (Phenomenex Luna, $C_8$(2), 250 X 10 mm, 5 µm) for purification using the following gradient solvent system: solvent A, water with 0.1% formic acid; solvent B, methanol with 0.1% formic acid; 10% B isocratic for 5 min, linear gradient to 80% B over 10 min, and then to 95% B over 60 min, with a flow rate of 2.5 mL/min. 15M-9E was eluted with minor impurities at a retention time of 50 min under these conditions. It was then subjected to another round of HPLC purification (Phenomenex Luna, $C_8$(2), 250 X 4.6 mm, 5 µm) using a modified gradient solvent system (10% B isocratic for 5 min, gradient to 85% B over 10 min, and then to 95% B over 30 min; flow rate, 0.8 mL/min). 15M-9E was isolated at a retention time of 34 min.

NMR spectroscopy and structural determination

All the NMR spectra were recorded using Bruker NEO 600 MHz NMR spectrometer. Chemical shifts are reported in ppm and referenced to the residual solvent signals of $CD_3OD$ ($\delta_H$ 3.30 for $^1H$ and $\delta_C$ 49.0 for $^{13}C$). The molecular formula of 15M-9E was determined as $C_{17}H_{32}O_2$ based on the HR-ESI-MS data ($[M–H]^-$ at *m/z* 267.2340, calculated for $C_{17}H_{31}O_2^-$, 267.2330). The complete chemical structure and one bond $^1H$–$^{13}C$ correlation assignments (Table S2) were established through a comprehensive analysis of 1D and 2D NMR data, including COSY, HSQC, and HMBC. Additionally, the position of the double bond was confirmed via GC-MS analysis.

High-resolution mass spectrometry for 15M-9E

High resolution mass spectrometry data were acquired using Agilent 1290 Infinity HPLC system coupled to an Agilent 6530 Quadrupole Time-of-Flight (QTOF) mass spectrometer equipped with an electrospray ionization (ESI) source operating in negative ion mode. The mass spectrometer was operated in full-scan mode with a mass scanning range of *m/z* 100-1700. A 5 µL aliquot of 15M-9E in methanol was injected onto a reversed-phase analytical column (Phenomenex Kinetex, $C_8$(2), 100 × 2.1 mm, 5 µm) using the following gradient solvent system: solvent A, water with 0.1% formic acid; solvent B, methanol with 0.1% formic acid; 10% B isocratic for 1 min, linear gradient to 100% B over 14 min. The gradient elution was performed at a flow rate of 0.4 mL/min. Data acquisition and analysis were performed using the Agilent MassHunter Qualitative Analysis software.

**Choanoflagellate genome editing and behavioral assays**

Choanoflagellate cell culture

A monoxenic culture *Salpingoeca rosetta* fed exclusively with the bacterium *Echinicola pacifica* (SrEpac, ATCC PRA-390) was used as the wild-type strain for all experiments and the basis for genome editing experiments. These cells were maintained in a synthetic sea water (Ricca 8363-5, filter-sterilized and referred to hereon as "SW") supplemented with 1.5% (v/v) of PYG (5 g/L peptone, 3 g/L yeast extract, 0.3% glycerol) and 1.5% (v/v) of CGM (cereal grass media from Fisher Science Education, 5 g/L, steeped for 3 hours in pre-boiled sea water and filtered). The media was additionally supplemented with L1 vitamins (Bigelow, 1:1000 dilution), L1 trace metals (Bigelow, 1:1000 dilution), L1 phosphate and nitrate (Bigelow, 1:1000), and 200 μM potassium iodide. Cells were maintained at 25 °C, typically split at a 1:5000 dilution every 2 days, and supplemented during each split with 1% of 10 mg/mL *E. pacifica*.

Genome editing

*Design and preparation of guide RNAs*

Genome editing followed previously published protocols for *Salpingoeca rosetta* (*19–21*). Candidate guide RNA sequences were obtained for each gene of interest using the EuPaGDT tool (http://grna.ctegd.uga.edu/) and the *S. rosetta* genome (Table S3). Guide RNA length was set at 15 and an expanded PAM consensus sequence, HNNRRVGGH, was used when possible. Genomic loci of interest were obtained from the Ensembl Protists hosting of the *S. rosetta* genome. Guide RNA candidates were filtered for guides with one on-target hit (including making sure the guides do not span exon-exon boundaries), zero off-target hits (including against the genome of the co-cultured bacterium *E. pacifica*) and lowest strength of the predicted secondary structure (assessed using the RNAfold web server: http://rna.tbi.univie.ac.at/cgi-bin/RNAWebSuite/RNAfold.cgi). crRNAs with the guide sequence of interest, as well as universal tracrRNAs, were ordered from IDT (Integrated DNA Technologies, Coralville, IA). Prior to the day of transfection, dried crRNA and tracrRNA from IDT were each resuspended in duplex buffer (30 mM HEPES-KOH pH 7.5; 100 mM potassium acetate, IDT Cat. No. 11-0103-01) to a concentration of 200 μM. Equal volumes of crRNA and tracrRNA were mixed, incubated for 5 minutes at 95 °C in an aluminum heating block, and then cooled to 25 °C slowly by removing the heat block from the heating source and cooling to RT. The annealed crRNA/tracrRNA is referred to as the gRNA and can be stored at -20 °C for weeks before use.

*Design and preparation of repair templates*

For the TRPW1 locus deletion, a double-stranded repair DNA template was generated by PCR. The base plasmid, pMS18 (Addgene #225681), encodes a puromycin resistance gene codon-optimized for *S. rosetta*, and flanked by *S. rosetta* regulatory elements to drive high expression. This cassette was amplified from pMS18 using homology arms to the TRPW1 locus. Two guide RNAs were chosen on either side of TRPW1 to fully excise the locus. The F primer for the template-generating PCR contained 50 bp of homology to the 5' side of the cut side for the N-terminal guide RNA, while the R primer contained 50 bp of homology to the 3' side of the C-terminal guide RNA. Between the genome annealing and plasmid annealing sequences was an 18-bp cassette – TTTATTTAATTAAATAAA – containing a stop codon in every possible reading frame (Table S3).

For the C-terminal ALFA-tagging of TRPW1, the pMS18 plasmid was digested with SacI and assembled with a gBlock (IDT) containing a short linker (GSG) and the ALFA peptide (SRLEEELRRRLTE) codon-optimized for *S. rosetta*. This is followed in the gBlock by a stop codon and the 3' untranslated region from the *S. rosetta* EFL gene (*22*). The assembly was done using NEB HiFi MasterMix according to the manufacturer's protocol. To generate the repair template for knock-in tagging, primers were designed to amplify the tag + the puromycin repair cassette from the plasmid with homology ends to either side of a gRNA cutting at the 3' end of the

TRPW1 locus. The amplification primers were chosen so that the puromycin cassette is inserted in the opposite orientation to the TRPW1 gene, i.e. the 3' ends of TRPW1 and $puro^R$ point towards one another.

Primers were ordered as Ultramers from IDT. To prepare the final repair DNA for CRISPR experiments, 100 µl PCRs were performed using 45x cycles, following by PCR purification (Qiagen) and concentration of the PCR product to ~4 µl volume on a 55 °C heat block. From this, 2 µl of concentrated PCR product were used for each 96-well transfection reaction.

*Transfection*

120 mL of choano culture was grown in a 3-layer flask (WestNet Inc Cat No 353143). To wash away bacteria from the choanoflagellates, the culture was split into three 50 ml conical tubes and centrifuged for 5 minutes at 2000 x g. The cell pellets were resuspended and combined in 50 ml of SW, followed by a 5 min spin at 2200 x g. The cells were washed once more with 50 ml SW and spun at 2400 x g. The pellet is resuspended in 100 µl SW and diluted 1:100 in SW for counting. Cells are diluted to $5 \times 10^7$ / mL in SW, then 100 µl aliquots (with $5 \times 10^6$ cells each) are prepared.

Priming buffer is prepared by diluting 10 µl of 1 mM papain (Sigma-Aldrich Cat. No. P3125-100MG) in 90 µl of dilution buffer (50 mM HEPES-KOH, pH 7.5, 200 mM NaCl, 20% glycerol, 10 mM cysteine). This is then diluted 1:100 in the rest of the priming buffer (40 mM HEPES-KOH, pH 7.5, 34 mM lithium citrate, 50 mM L-cysteine, 15% PEG-8000) for a final concentration of 1 µM papain. Equal volumes of pre-annealed gRNA and *Sp*Cas9 (20 µM, NEB Cat. No. M0646M) are mixed and incubated for 1 hour at RT to form the RNP. 4 µl of RNP is used per transfection reaction.

Each aliquot of cells is spun at 800 x g for 5 minutes and resuspended in 100 µl priming buffer and incubated for 35 minutes at RT. The priming reaction is quenched by adding 10 µl of 50 mg/ml bovine serum albumin fraction V (Thermo Fisher Scientific Cat. No. BP1600-1000). Cells are spun at 1250 x g for 5 minutes and resuspended in 50 µl Lonza SF buffer (Lonza Cat. No. V4SC-2960). For each transfection, 16 µl of Lonza SF buffer is mixed with 4 µl of RNP targeting the gene of interest, 2 µl of repair oligo (previously concentrated PCR product), and 1 µl of washed and primed cells.

The nucleofection reactions are added to a 96-well nucleofection plate (Lonza Cat. No. V4SC-2960) and pulsed with a DG137 pulse (*21*) in the Lonza 4D-Nucleofector (Cat. No. AAF-1003B for the core unit and AAF-1003S for the 96-well unit).

After the pulse, 100 µl of ice-cold recovery buffer (10 mM HEPES-KOH, pH 7.5; 0.9 M sorbitol; 8% [wt/vol] PEG 8000) is immediately added to each well of the nucleofection plate and incubated for 5 minutes. Then the entire contents of the well are added to 2 mL of choanoflagellate media with 1% (v/v) of a 10 mg/mL *E. pacifica* suspension in a 6-well plate and cultured at 25 °C.

*Selection and genotyping*

24 hours after transfection, 80 µg/mL of puromycin (Gibco) was added to choanoflagellate cultures. This led to most cells dying, with resistant cells re-populating the well after 3-4 days of antibiotic selection. To isolate clonal populations, resistant cells were diluted to 1 cell/mL in choano media supplemented with 1% of 10 mg/mL *E. pacifica* and 80 µg/mL puromycin and plated at 200 µl/well in 96-well plates. After 3-4 days of growth, clonal populations were identified for DNA extraction and genotyping PCRs. DNA extraction was done using NucleoType

DirectLyse reagent according to manufacturer's protocol, using 100 µl of dense choano culture as input. Genotyping PCRs were performed using Q5 DNA polymerase supplemented with 1M betaine (see Table S3 for primer sequences). PCR products were analyzed by gel electrophoresis and submitted for long-read amplicon sequencing (Azenta) to confirm genomic changes.

Choanoflagellate accumulation at *Algoriphagus machipongonensis* biofilms

*Algoriphagus machipongonensis* was grown in liquid culture in PYG media. *S. rosetta* was grown to mid-log phase in standard media with *E. pacifica*. Microfluidic chambers were built between two glass coverslips with glass capillaries as spacers, and choanoflagellate culture was added between the glass coverslips. *A. mac* biofilm was scraped from the wall of a liquid culture with another glass capillary and positioned in the center of the chamber. Images were recorded every 10 seconds using an Olympus IX73 inverted microscope with a 10X UPlanSApo objective (0.40 NA, Olympus), a TF4-100 light source (Olympus) and an ORCA-Flash 4.0 camera (Hamamatsu), controlled using MetaMorph software. For presentation in Figure 1A, the edge of the bacterial biofilm is labeled with a dotted line and individual choano cells are labeled with white dots.

Choanoflagellate surface accumulation assays

To prepare embedded drug pucks, a 2% molten agarose solution in SW was prepared. When the agarose cooled to 75-80 °C, 100 µL of molten agarose was mixed with 1 µL of drug solution or vehicle control using a cut P200 tip and deposited on a petri dish, then immediately covered with a 22 x 30 mm glass coverslip, allowing the agarose to spread underneath to a consistent thickness. After 5 minutes, the coverslip was turned over, and a folded wet paper towel was placed in the dish to prevent the agarose from drying out. 2 mm circles were cut out using a biopsy punch (World Precision Instruments Cat No 504531). For RS 39604, 1 µL of a 50 mM stock (in DMSO) was used to generate a final concentration of 500 µM in the agarose puck, which is on the edge of its solubility in sea water, leading to some visible precipitate. For vehicle control, 1 µL of DMSO was used instead. 1 mL of *S. rosetta* cells at mid-log density (4e5 to 9e5 cells/mL) was stained with 1 µL of CellTrace CFSE dye (Thermo Fisher Cat No C34570) from a stock concentration of 5 mM in DMSO, vortexed briefly and spun at 3000 x g for 7 minutes. While spinning, an agar puck was transferred with tweezers to an 8-well ibiTreat µ-Slide (Ibidi Cat No 80826). The µ-Slide was also kept in a petri dish with a wet, folded paper towel to keep high humidity and prevent the agarose puck from drying out. After spinning, cells were resuspended in 200 µL SW. At the microscope, 250 µL of SW were added to the well with the agarose puck and the stage positioned so that the convex edge of the puck protrudes about 1/3 into the field-of-view of the 10X objective. 10 µL of resuspended cells were added and imaged every 2 seconds for the duration of the time course. Image acquisition was done using the previously described Olympus IX73 set-up, using a Lambda 421 (Sutter) illumination system for fluorescence imaging.

For quantification, a square ROI was chosen (1024 x 1024 px) corresponding to the left half of the 2048 x 2048 FOV, centered between top and bottom. In FIJI, images were converted to 8-bit and background subtracted with a 25 px rolling ball method. Particles were identified and counted using the FIJI "Find maxima" function, adjusting the prominence threshold manually for each image stack, verifying correct cell assignments at various points of the video. Prominence thresholds ranged from 80 to 150. For representative images in the figures, the outline of the agar puck was drawn using a brightfield image collected at the beginning of the time course, and cells were displayed by thresholding the fluorescent image and using the binary dilate function in FIJI 8 consecutive times.

Choanoflagellate swimming behavior assays

To analyze swimming behavior in the presence of 25 μM 15M-9E, cells were grown to mid-log phase, counted, and diluted to 50,000 – 100,000 cells/mL in SW. 250 μL of this cell suspension was mixed with 0.3 μL of 15M-9E stock solution (5 mg/mL, or 18.7 mM) and added to an 8-well ibiTreat μ-Slide (Ibidi Cat No 80826). Cells were allowed to settle for exactly 1 min before image acquisition. Image acquisition was done using the previously described Olympus IX73 set-up, imaging at a theoretical speed of 10 frames per second. Because the actual imaging speed was slower than this and slightly variable experiment to experiment (always between 6 and 7 fps), the actual fps was calculated from the experiment metadata and used to normalize swimming track velocities. Videos lasted from 89 to 102 seconds, depending on camera speed. Videos were acquired with the cells slightly out of focus so that they approximated black circles, which aided in downstream tracking and analysis.

For analysis in FIJI, videos were thresholded and TrackMate was used with the mask detector option after dilating the thresholded images 2-3 times. Spots were filtered for circularity > 0.50 and area > 30 $px^2$. The LAP tracker algorithm was used for connecting spots, with 30.0 px frame-to-frame linking, 30.0 px gap closing, a 5-frame max gap, and no splitting or merging allowed. Tracks were filtered for those lasting >= 150 frames. Track statistics were calculated by TrackMate. Mean track speed in px/frame was converted to μm/s using a 0.82 μm/px conversion factor and the video fps calculated from the image metadata. For plotting, individual tracks are shown in grey, but for each experiment the mean track speeds for all tracks are combined into a weighted average using the duration of the tracks as the weighting factor, which is displayed for each biological replicate as a larger dot. This prevents over-counting cells which had a trajectory that was split into multiple shorter tracks.

High-speed flagellar imaging

Choanoflagellates were grown to mid-log phase, and washed twice in synthetic SW, spinning for 5 min at 3000 x to pellet cells. 50 μL of cells were injected into a microfluidic imaging slide (μ-Slide VI, poly-L-lysine surface, Ibidi Cat. No. 80604) and settled for 15 minutes. Using a Zeiss ELYRA microscope with a 100X Plan-APOCHROMAT oil immersion objective (1.40 NA, Zeiss), an HXP120V light source, and an ORCA Fusion BT camera (Hamamatsu). Videos were acquired at 433 frames/second, totaling 52,000 frames for a 2-minute video. Immediately after starting the video, 100 μL of SW or SW + drug was added to the inlet port of the microfluidic channel. Flows subsided after 5-10 seconds. For each data point, the same exact cell was recorded first with the addition of SW, and then with the addition of drug. All trials with 15M-9E utilized a 75 μM concentration. Between SW and drug additions, 100 μL was removed from the outlet port. If the cell lost attachment to the surface in the field-of-view during any step of the imaging, it was excluded from further analysis.

To extract flagellar beating data from videos, a computational pipeline was combined with manual annotation and supervision. First, custom FIJI macros were used to sub-sample the videos, prompt the user to draw line ROIs, and extract average intensity across the line ROI for the duration of the video. A custom Python script was used to extract flagellar frequency and beat power from the intensity values and set a threshold for beat power, below which the flagellum was labeled as paralyzed. The output graph of paralyzed vs. beating periods was compared with a manual inspection of the sub-sampled video. Brief (<1 s) false positive paralysis episodes (from comparing manual inspection to script output) were tolerated, but any greater discordance between the output annotations and the visual inspection of the data prompted us to re-draw line ROIs and re-analyze intensity profiles and flagellar beating over time, until agreement was reached between output annotations and manual inspections.

The Python script median-centered each trace and bandpass-filtered each trace from 10-80 Hz, using a third-order Butterworth bandpass filter. The script detected periodic peaks and troughs, retaining one event train (peaks or troughs) based on event number and regularity of spacing. Candidate beats were adaptively pruned when multiple peaks occurred closer together than the recent local inter-peak interval, retaining the stronger beat. A smoothed beat frequency was calculated over a sliding window of 10 cycles. For display in Figure 3D, an additional smoothing over a 20-cycle sliding window was also calculated. Beat "power" was calculated as the sliding window (0.5s) root-mean-square amplitude of the bandpass-filtered signal after local median subtraction. To set a threshold of beat power for calling paralysis vs. active beating, a reference power was established by taking the median of the top 20% of the beat power distribution, then setting the paralysis threshold as a fraction of that reference power, default 30%. Paralysis intervals were called as contiguous periods during which the beat power fell below the threshold, lasting at least 0.5s, and with a sufficiently steep onset, by computing the minimum slope of beat power within the last 0.35s and comparing this to a defined threshold, set by a "slope factor" defaulting to 0.35. For reporting of paralysis episodes, the first 10s of the recording were excluded due to both flagellar paralysis and changes in cell orientation caused by the onset of flow into the microfluidic chamber. Appropriate parameter values were ascertained by iteratively comparing script predictions to manual observations of flagellar beating.

For flagellar imaging of rosettes, *S. rosetta* in mid-log growth phase was induced with outer membrane vesicles (OMVs) from *Algoriphagus machipongonensis* (see above for preparation protocol). The OMV dilution used to get robust induction (~100% of cells in rosettes) was determined empirically for each batch, ranging from 1:100 to 1:500. Because it is difficult to get rosettes to stick stably on poly-lysine coated surfaces, rosettes were used for experiments 6-10 hours after induction, when they consist of between 2 and 4 cells. Rosettes still frequently lost attachment to the surface of the microfluidic channel during agonist addition. Because of this, rosette flagellar analysis was performed with the commercially available synthetic agonist RS 39604 (at 50-100 μM) and not 15M-9E, which requires labor-intensive purification. For quantification of rosette coordination, the first paralysis event after flow had subsided from agonist addition was analyzed, with coordination called when multiple cells arrested within 100 ms of each other. In most instances, the gap was much shorter but occasionally one cell completed a very slow beat cycle before complete arrest, lasting up to 100 ms, so this was still included as "coordinated". Three biological replicates were performed with n = 5, 4, and 6 rosettes in each set.

During high-speed imaging, we observed that extended exposure to TRPW1 agonists also led to other phenotypes. For 15M-9E, initial bouts of paralysis were sometimes followed by microvilli thickening and rarely by cell cortex contractions. For RS 39604, initial bouts of paralysis were sometimes followed by cell cortex contractions, and rarely by flagellar disassembly, microvilli disassembly, and cell death. We attribute these responses to persistent calcium influx into the cell through TRPW1, and perhaps "off-target" effects on the 16 other TRPW homologs or other TRP channels in *S. rosetta*. In the natural environment, where TRPW1 most likely detects bacterial lipids through contact-dependent striking of the flagellum against bacterial membranes or outer membrane vesicles, compared to the whole cell being bathed in agonist, these other phenotypes are likely limited.

Phagocytosis of polystyrene beads

Wild-type and TRPW1$^{KO}$ cells were grown to mid-log phase. 250 μL of cell suspension was added to each well of an Ibidi 8-well poly-L-lysine coated dish (Ibidi Cat No 80824). Cells were left to attach for 15 minutes, while 1 μL of 1 μm polystyrene beads (Thermo Fisher Cat No F8851) was mixed with 1 mL of synthetic sea water and bath sonicated for 15 minutes to disperse bead clumps. After cell settling, 200 μL of media was replaced with 200 μL of the bead suspension. Cells were incubated with beads for 2.5 hours before imaging on a Zeiss LSM 880 with a 63X Plan-APOCHROMAT oil immersion objective (Zeiss). For each condition, 3 x 2 tiles were collected, using a 10 μm stack with 1.0 μm slices. Numbers of beads per cell was annotated manually.

Growth curve

Wild-type and TRPW1$^{KO}$ cells were grown to mid-log phase and counted using a LUNA-FL automatic cell counter. For each genotype, cells were diluted to 500 cells/mL, supplementing with 1% (v/v) of a 10 mg/mL *E. pacifica* food pellet. 500 μL/well of diluted cells were added to a 24-well plate and cultured at 22 °C. Approximately every 12 hours (exact timing is shown in the data points of the growth curve itself), three replicates per genotype were fixed with 10 μL of 32% paraformaldehyde. Cell counts were determined using a LUNA-FL automatic cell counter.

Immunofluorescence

Wells in a 96-well imaging plate (μ-Plate, ibiTreat, Ibidi Cat No 89626) were washed twice with 100% ethanol, twice with water, and incubated with 100 μL of 0.1 mg/mL poly-D-lysine (Gibco) for 15 minutes, then washed 3 times with water. Cells (wild-type and TRPW1$^{ALFA}$) were grown to mid-log phase and 100 μL of cell culture was added to the coated wells and incubated for 20 minutes to allow cell attachment. The cells were washed twice with 200 μL of 4X PEM buffer (400 mM PIPES, 4 mM EGTA, 4 mM $MgCl_2$), using gel loading tips and slow, careful addition and removal of liquid. After the second wash, only 100 μL was removed, leaving 200 μL volume. To this was added 200 μL of freshly prepared fixative (3 parts 4X PEM to 1 part 32% paraformaldehyde), and fixation proceeded for 15 minutes at RT. After fixation, 300 μL was removed, and wells were washed 3x with 200 μL of 4X PEM, leaving only 100 μL at the end. Blocking/permeabilization was done with the addition of 200 μL of Intercept PBS Blocking Buffer (Licor) + 1% Triton X-100, incubating for 30 minutes at RT. After this, 200 μL was removed, and 100 μL of nanobody solution was added: Intercept PBS + 1% Triton X-100 1:100 dilution of FluoTag X2 anti-ALFA Alexa647 nanobody (NanoTag Biotechnologies Cat. No. N1502-AF647-L). The nanobody was incubated overnight at 4 °C. The following day, 100 μL of volume was removed, and wells were washed 3x with 200 μL of 4X PEM, leaving 100 μL volume remaining. Finally, 100 μL of phalloidin 488 (AAT Bioquest Cat. No. 23115) diluted 1:1000 in 4X PEM was added to each well and incubated for 30 mins at RT before imaging. Imaging was done on a Nikon Eclipse Ti2 with a CSU-W1 SoRa spinning disk (Yokogawa).

Quantification of fluorescent signal along flagellum

For TRPW1$^{ALFA}$ images, a maximum intensity projection of the z-stack was performed, followed by manual tracing of individual flagella using a segmented line ROI in FIJI. Intensity profiles for each flagellum were extracted. Data from multiple flagella was normalized and interpolated along flagellar length, with intensity values normalized to the highest intensity pixel as 1. The average of all intensity profiles was plotted in Prism 10 using standard error of the mean as a bounding box. 20 flagella with visible TRPW1$^{ALFA}$ signal in the flagellum were selected for tracing and quantification.

Western blot

Cells were grown to mid-log phase and 50 mL of culture was concentrated in 1 mL of SW. An aliquot of 10 x $10^6$ cells was made, spun for 5 mins at 6000 x g, and resuspended in 100 μL of a digitonin lysis buffer (10 mM digitonin, 20 mM Tris pH 8.0, 150 mM KCl, 5 mM $MgCl_2$, 250 mM sucrose, 1 mM Pefabloc, 1 Roche cOmplete mini protease inhibitor tablet/5 mL). Right before lysis, 1 mL of base lysis buffer was supplemented with 1 μL of 1 M DTT and 15 μL of Pierce Universal Nuclease (Thermo Fisher Scientific Cat. No. 88701). Cells were lysed for 30 mins at 4 °C, and the lysate was clarified by spinning at 6000 x g for 10 mins at 4 °C. The supernatant was removed and combined with 10 μL of 10% Tween-20 and incubated for 10 mins at RT. 20 μL of sample was mixed with 20 μL of 2X Laemmli Sample Buffer supplemented with β-mercaptoethanol (190 μL of 2X Laemmli + 10 μL of βME).

20 μL of samples were added to a Bio-Rad Mini-PROTEAN TGX Precast Protein Gel 4-20% (BioRad Cat No 45621093) and gel electrophoresis was done for 40 minutes at 200V. Transfer to nitrocellulose membrane was performed using a Trans-Blot Turbo machine (BioRad) with a nitrocellulose transfer pack (BioRad Cat. No. 1704158), running at 25V for 30 m. Membranes were blocked in Intercept PBS Blocking Buffer (LICOR) for 1 hour at RT. ALFA-tagged proteins were detected with a FluoTag-X2 anti-ALFA LICOR IRDye 800CW nanobody (NanoTag Biotechnologies N1502-Li800-L), at a 1:10,000 dilution in Intercept PBS, incubating for 2 hours at RT. The membrane was washed 3x for 5 mins with PBST (0.2% Tween-20 in PBS), then rinsed with PBS and imaged on a LICOR Odyssey.

**Structural biology**

Protein expression and purification

For cryo-EM studies, TRPW1 bacmids and baculoviruses were produced using standard procedures (*18*). Briefly, baculovirus was made in Sf9 cells (Thermo Fisher Scientific, mycoplasma test negative, GIBCO #12659017) for 96-120 hours and added to suspension-adapted HEK 293S cells lacking N-acetyl-glucosaminyltransferase I (GnTI$^-$, mycoplasma test negative, ATCC #CRL-3022) that were maintained at 37°C and 6% $CO_2$ in Freestyle 293 media (Gibco-Life Technologies #12338-018) supplemented with 2% FBS. To enhance protein expression, sodium butyrate (10 mM) was added 24 hours after transduction, and the temperature was reduced to 30 °C. The cells were harvested by 15-min centrifugation at 5,471 g using a Sorvall Evolution RC centrifuge (Thermo Fisher Scientific) 72 hours after transduction. The cells were washed in the phosphate buffer saline (PBS) pH 8.0 and pelleted by centrifugation at 3,202 g for 10 min using an Eppendorf 5810 centrifuge.

The cell pellets for each construct were resuspended in the ice-cold buffer containing 20 mM Tris pH 8.0, 150 mM NaCl, 0.8 μM aprotinin, 4.3 μM leupeptin, 2 μM pepstatin A, 1 μM phenylmethylsulfonyl fluoride (PMSF), and 1 mM β-mercaptoethanol (βME). The suspension was supplemented with 1% (w/v) digitonin, and cells were lysed at constant stirring for 2 hours at 4°C. Insoluble material was removed by ultracentrifugation for 1 hour at 186,000 g in a Beckman Coulter centrifuge using a 45 Ti rotor. The supernatant was added to the strep resin, which was then rotated for 30 min at 4 °C. The resin was washed with 20 column volumes of wash buffer containing 20 mM Tris pH 8.0, 150 mM NaCl, 1 mM βME, and 0.06% (w/v) digitonin, and the protein was eluted with the same buffer supplemented with 2.5 mM D-desthiobiotin. The eluted protein was concentrated to 0.5 ml using a 100-kDa NMWL centrifugal filter (MilliporeSigma Amicon) and then centrifuged in a Sorvall MTX 150 Micro-Ultracentrifuge (Thermo Fisher Scientific) for 30 min at 186,000 g and 4°C using a S100AT4 rotor before injecting it into a size-exclusion chromatography (SEC) column. The protein was purified using a Superose 6 10/300 GL SEC column attached to an AKTA FPLC (GE Healthcare) and equilibrated with the buffer containing 20 mM Tris pH 8.0, 150 mM NaCl, 1 mM βME, and 0.06% (w/v) digitonin. The tetrameric peak fractions were pooled and concentrated to 4.0-4.5 mg/ml using a 100-kDa NMWL centrifugal filter (MilliporeSigma Amicon).

Cryo-EM sample preparation and data collection

Prior to sample application, UltrAuFoil R 1.2/1.3 (Au300) grids were plasma treated in a PELCO easiGlow glow discharge cleaning system (0.39 mBar, 15 mA, "glow" for 25 s, and "hold" for 10 s). A Mark IV Vitrobot (Thermo Fisher Scientific) set to 100% humidity at 4 °C was used to plunge-freeze the grids in liquid ethane after applying 3 μl of protein sample to their gold-coated side using the blot time of 5 s, blot force of 5, and wait time of 15 s. The grids were stored in liquid nitrogen before imaging.

Images of frozen-hydrated particles of TRPW1 were collected at the Columbia University Cryo-EM Facility using the Leginon software (*23*) on a Titan Krios transmission electron microscope (TEM) (Thermo Fisher Scientific) operating at 300 kV and equipped with a post-column GIF Quantum energy filter and a Gatan K3 Summit direct electron detection (DED) camera (Gatan, Pleasanton, CA, USA). The total of 5,236 and 3,651 micrographs were collected for TRPW1 in the apo and pre-active state bound to RS 39604, respectively. Micrographs were collected in the counting mode with an image pixel size of 0.825 Å across the defocus range of −0.5 to −2.5 μm. The total dose of ~58 $e^{-}Å^{-2}$ was attained by using the dose rate of ~16 $e^{-}pixel^{-1}s^{-1}$ across 50 frames during the 2.5-s exposure time.

Image processing and 3D reconstruction

Data were processed in cryoSPARC v4.5.3 (*24*). Movie frames were aligned using the Patch Motion Correction algorithm implemented in cryoSPARC. After contrast transfer function (CTF) estimation, micrographs were manually inspected to remove those with lower predicted CTF-correlated resolution and outliers in defocus values, ice thickness, and astigmatism. Similar processing workflow was used for both datasets.

For example, for TRPW1 in the apo state, the total number of 944,482 particles were picked using internally-generated 2D templates and extracted with 512-pixel box size and then binned to 256-pixel size. After several rounds of reference-free 2D classification and heterogeneous refinement with one reference class and three automatically generated "garbage" classes, the best 245,802 "consensus" particles were re-extracted with 400-pixel box size without binning and then subjected to reference-based motion correction in cryoSPARC. These particles were 3D classified into 5 classes, resulting in one well-defined class in the apo conformation. The particles corresponding to this well-defined class were subjected to the final non-uniform refinement, with C4 rotational symmetry imposed, and yielded a 2.37-Å resolution 3D reconstruction. To improve the quality of reconstruction for the beta-barrel domain that experienced rigid-body movements, the final set of particles was C4 symmetry expanded (824,888 particles) and subjected to the local refinement with a mask around the beta-barrel, hinge, and linker domains, yielding a map with the resolution of 2.57 Å. The reported resolution for the final maps was estimated in cryoSPARC using the gold standard Fourier shell correlation (GSFSC) using the FSC = 0.143 criterion. For model building, a composite map was also created from the maps for the entire channel and the beta-barrel domain using the Combine-focused-maps package implemented in Phenix. The local resolution was calculated in cryoSPARC using the FSC = 0.5 criterion. Cryo-EM densities were visualized using UCSF ChimeraX (*25*). For structural comparisons, the following PDB structures were download: TRPV1 (2PNN), TRPM4 (5WP6), TRPM2 (6DRJ), TRPV1-capsaicin (7LR0).

**Molecular dynamics**

Simulations

The structural model of the full-length TRPW1 channel in the RS39604-bound state (TRPW1-RS 39604), containing four $Ca^{2+}$ ions resolved at the S2–S3 calcium-binding sites in the cryo-EM structure, was inserted into a hydrated lipid bilayer consisting of palmitoyloleoylphosphatidylcholine (POPC) lipids (~600 lipids, ~160 × 160 × 155 $Å^3$ modeling cell size). Four molecules of 15-methylhexadec-9-enoic acid (15M-9E) were placed simultaneously into the putative binding sites, located within the transmembrane domain at the interface between the S4–S5 linker, S1–S4 bundle, and TRP helix. $Na^+$ and $Cl^-$ ions were added to achieve physiological salt concentration (150 mM) and electroneutrality. Three independent simulation replicas were prepared.

All replicas were equilibrated in several consecutive stages: $5 \times 10^4$ steps of steepest descent energy minimization, followed by heating from 5 to 310 K during a 200-ps MD run, then 10 ns of MD with fixed positions of all protein heavy atoms, 10 ns of MD with fixed positions of the protein backbone, and 50 ns of MD with fixed positions of the protein Cα atoms to allow membrane relaxation. The force constant k = 10 kJ/(mol × $Å^2$) was applied for all positional restraints during equilibration. The 15M-9E molecules were free to move during all equilibration stages. Production MD runs of 500 ns were then carried out for each replica.

MD simulations were performed using GROMACS 2024.4 (*26*), CHARMM36m force field (*27*) with NBFIX corrections (*28*), and the TIP3P water model (*29*). Simulations were carried out with an integration time step of 2 fs; hydrogen-containing bond lengths were constrained using the LINCS algorithm (*30*). 3D periodic boundary conditions were imposed. Constant temperature of 310 K was maintained by the v-rescale thermostat (*31*), and constant semi-isotropic pressure of 1 bar was maintained by the Parrinello-Rahman barostat (*32*). A cutoff distance of 12 Å was applied for evaluation of nonbonded interactions, and the particle-mesh Ewald (PME) method (*33*) was employed for treatment of long-range electrostatics. The topology and partial atomic charges of 15M-9E were generated using the CHARMM General Force Field (CGenFF) (*34*, *35*).

Analysis

The physicochemical properties of the TRPW1 ion-conducting pore were characterized using the dynamic molecular portrait (DMP) approach (*36*). Cylindrical projections of the molecular hydrophobicity potential (MHP) were calculated on the pore surface to map the distribution of hydrophobic and hydrophilic regions along the pore axis. MHP values were computed using atomic hydrophobicity constants according to Wildman and Crippen (*37*), and visualized using the Molecular Surface Topography (MST) tool available at https://model.nmr.ru/cell (*36*).

Hydrogen bonds (HB) and salt bridges (SB) between 15M-9E and TRPW1 were identified using standard geometric criteria: an HB was considered present when the distance between the donor (D) and acceptor (A) heavy atoms was less than 3.5 Å and the D–H···A angle was 180 ± 30°; a

SB was defined as an interaction between oppositely charged groups with a D–A distance below 4.0 Å. Bond occupancy was defined as the fraction of MD trajectory frames in which a given contact was present. All intermolecular interactions were calculated using the Contacts tool (https://model.nmr.ru/cell). MD data were visualized using PyMOL (*38*).

### Supplementary Figures

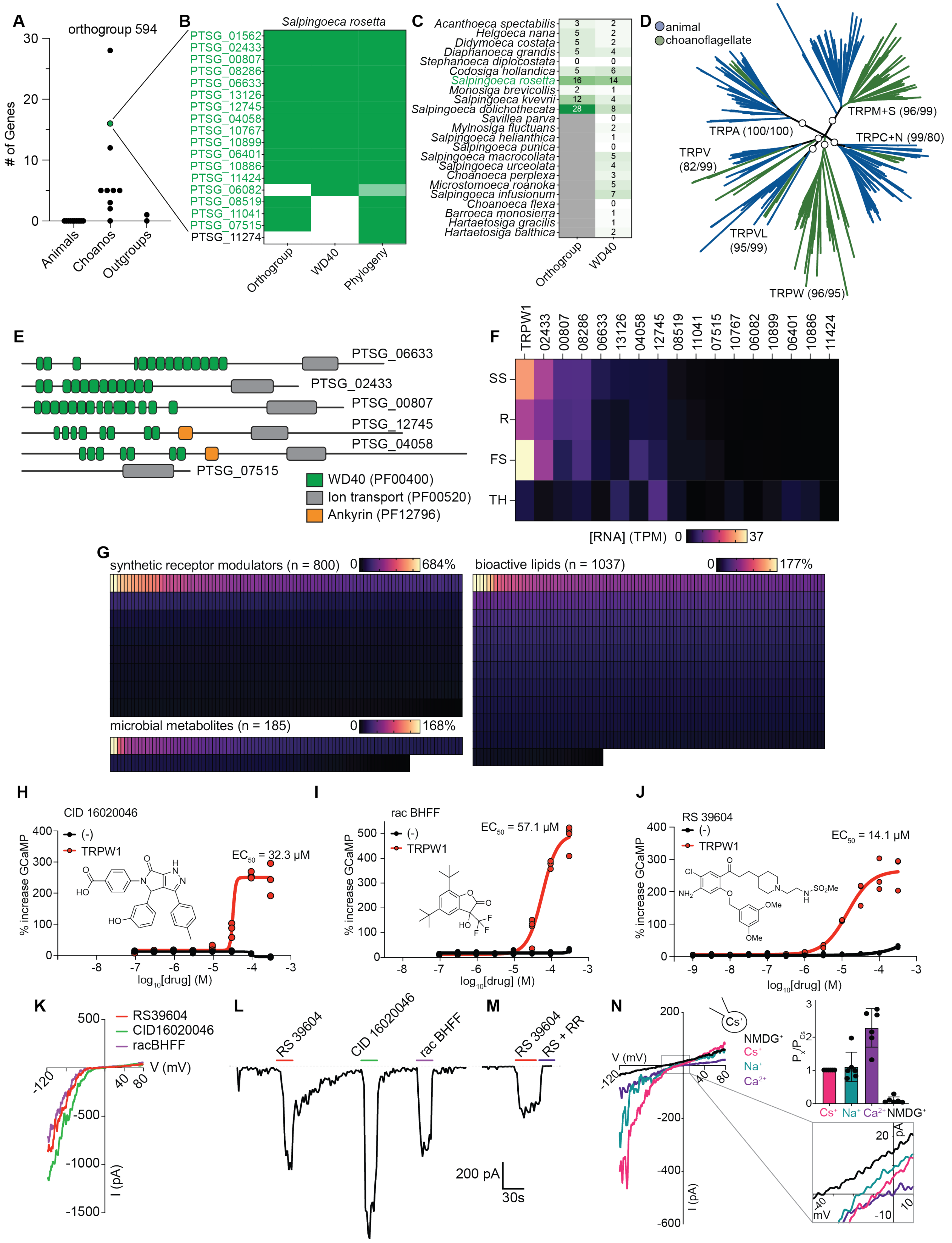

**Figure S1. Identification of an ion channel receptor family in choanoflagellates.** (**A**) An OrthoFinder analysis of diverse animal, choanoflagellate, and outgroup proteomes identified orthogroup number 594 that was present in an outgroup, expanded in choanoflagellates, and absent in animals. Inspection of predicted domains in this orthogroup identified WD40 repeats (PF0400) and an ion channel domain (PF00520). (**B**) The full set of TRPW family members in *S. rosetta*. The original orthogroup predicted 16 members, 13 of which contained WD40 repeats. A phylogenetic analysis (Figure 1B) of all related genes in the voltage-gated channel superfamily revealed one more TRPW, PTSG_06082, which contained WD40 repeats and branched as part of a sister clade to the other TRPWs that included this sequence and the *S. rosetta* TRPV homolog. Phylogenetic analysis also included one other gene, PTSG_11274, in the TRPW branch, but based on domain architecture, we assign this gene as a TRPM family member (see Materials and Methods for more detail). (**C**) TRPWs are found across the choanoflagellate clade. Reported here are numbers of TRPW family members by the original OrthoFinder analysis and by a search for the WD40 + ion channel architecture across all available choanoflagellate proteomes on EukProt. (**D**) TRPWs form a distinct sub-family among type I TRP channels. For display, all branches with <70% bootstrap support by UFboot are collapsed into palintomies. Circles indicate node support for the various TRP clades, with values indicating UFboot/SH-aLRT values. (**E**) Diverse domain architectures of TRPW family members in *S. rosetta*. (**F**) Expression of TRPW family members across *S. rosetta* life history stages by RNA sequencing. SS = slow swimmer, the free-swimming growth stage of this species in the absence of rosette-inducing bacteria. R = rosettes. FS = fast swimmers, a cell state induced by starvation. TH = thecate, a cell type with an extracellular anchoring to an environmental substrate. RNA sequencing analysis was done as reported by Coyle *et. al.* (*39*), with the full dataset available in Leon *et. al.* (*40*). (**G**) Screening results for *S. rosetta* TRPW1 heterologously expressed in HEK293Tcells with 10 μM of each molecule from the following commercially available libraries: synthetic receptor modulators (Tocris Cat. No. 7152, 800 molecules), bioactive lipids (Cayman Cat. No 10506, 1037 molecules), and microbial metabolites (Millipore Sigma GUTMLS, 185 molecules). (**H**) Dose-response relationship of TRPW1 heterologously expressed in HEK293Tcells against CID 16020046. n = 3 $Ca^{2+}$ imaging replicates per concentration. $EC_{50}$ = 32.3 μM. (**I**) Dose-response relationship of TRPW1 heterologously expressed in HEK293Tcells against rac BHFF. n = 3 $Ca^{2+}$ imaging replicates per concentration. $EC_{50}$ = 57.1 μM (95% CI from 49.5 μM to 65.8 μM). (**J**) Dose-response relationship of TRPW1 heterologously expressed in HEK293Tcells against RS 39604, with half-log dilutions from 300 μM to 1 nM. n = 3 $Ca^{2+}$ imaging replicates per concentration. $EC_{50}$ = 14.1 μM (95% CI from 10.6 μM to 19.7 μM). (**K**) Current-voltage relationships of TRPW1 heterologously expressed in HEK293Tcells in response to 30 μM RS 39604, 100 μM CID 16020046, 100 μM rac BHFF. Representative of n = 5 cells. (**L**) Representative kinetic trace of whole-cell currents evoked by the listed TRPW1 agonists. Representative of n = 5 cells. Currents measured in response to -120 to 80 mV voltage ramps and current measured at -120 mV plotted over time. (**M**) Representative current evoked by 100 μM RS 39604 was by 10 μM ruthenium red (RR). Representative of n = 8 cells. (**N**) Voltage-current relationships of TRPW1 heterologously expressed in HEK293Tcells for cation substitution experiments. Representative of n = 6 cells. The average $E_{rev}$ for each cation was used to calculate relative permeabilities using the Goldman-Hodgkin-Katz equation (see Materials and Methods). Average +/- S.D. for n = 6 cells. 95% CI for $P_{Ca}/P_{Cs}$: 1.67 – 2.89.

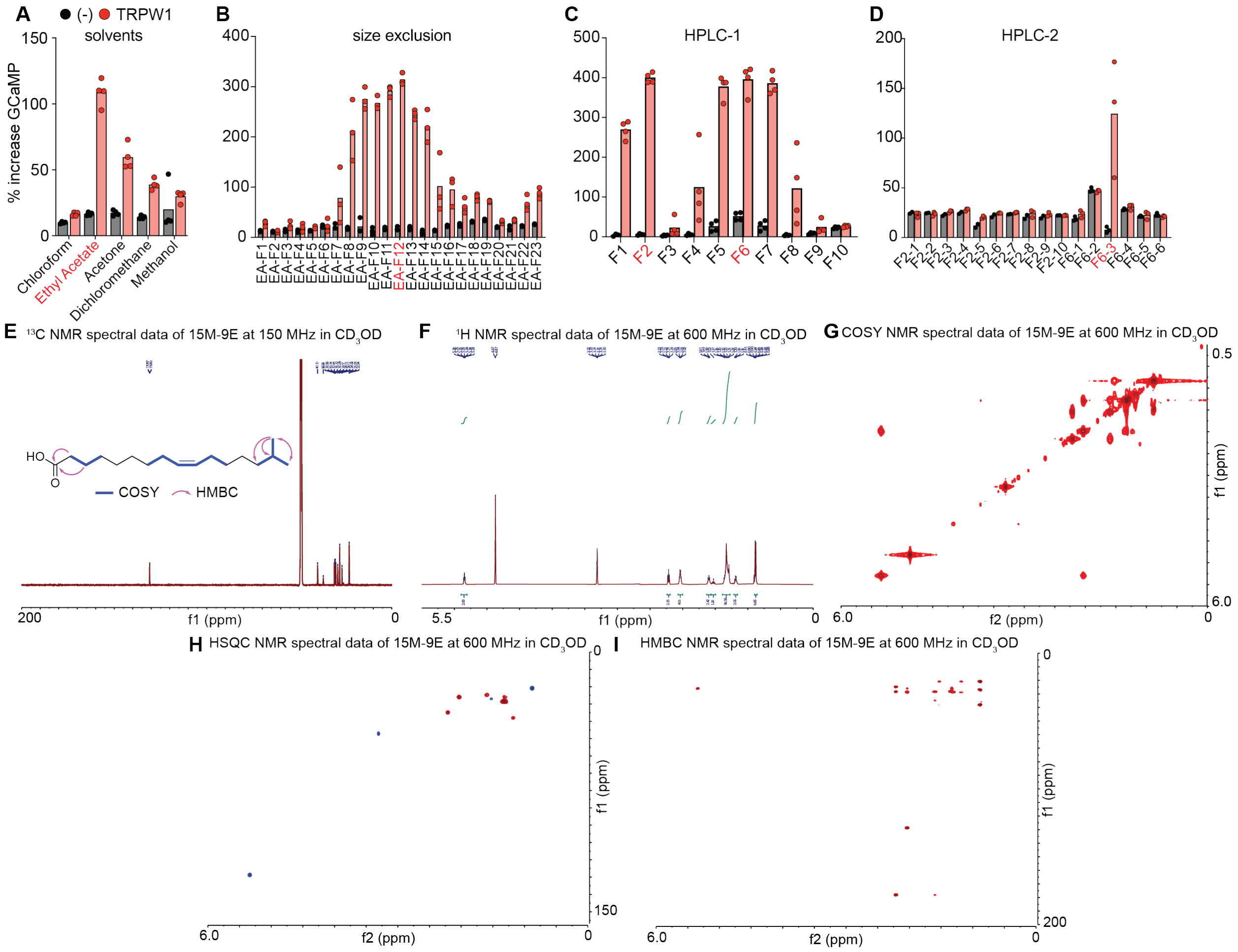


**Figure S2. Fractionation of *A. mac* extract and small molecule agonist identification.** (**A**) Activity of extracts from *Algoriphagus machipongonensis* in different solvents against TRPW1 heterologously expressed in HEK293Tcells. n = 4 $Ca^{2+}$ imaging replicates per fraction. (**B**) Activity of size exclusion column fractions from *Algoriphagus machipongonensis* against TRPW1 heterologously expressed in HEK293Tcells. n = 3 $Ca^{2+}$ imaging replicates per fraction. (**C**) Activity of HPLC C8 column fractions from *Algoriphagus machipongonensis* against TRPW1 heterologously expressed in HEK293Tcells. n = 4 $Ca^{2+}$ imaging replicates per fraction. (**D**) Activity of a second round of HPLC C8 column fractionation from *Algoriphagus machipongonensis* against TRPW1 heterologously expressed in HEK293. n = 3 $Ca^{2+}$ imaging replicates per fraction. (**E**) $^{13}$C NMR spectral data of 15M-9E at 150 MHz in CD3OD. See Materials and Methods for further detail on molecular identification. (**F**) $^1$H NMR spectral data of 15M-9E at 600 MHz in CD3OD. (**G**) COSY NMR spectral data of 15M-9E at 600 MHz in CD3OD. (**H**) HSQC NMR spectral data of 15M-9E at 600 MHz in CD3OD. (**I**) HMBC NMR spectral data of 15M-9E at 600 MHz in CD3OD.

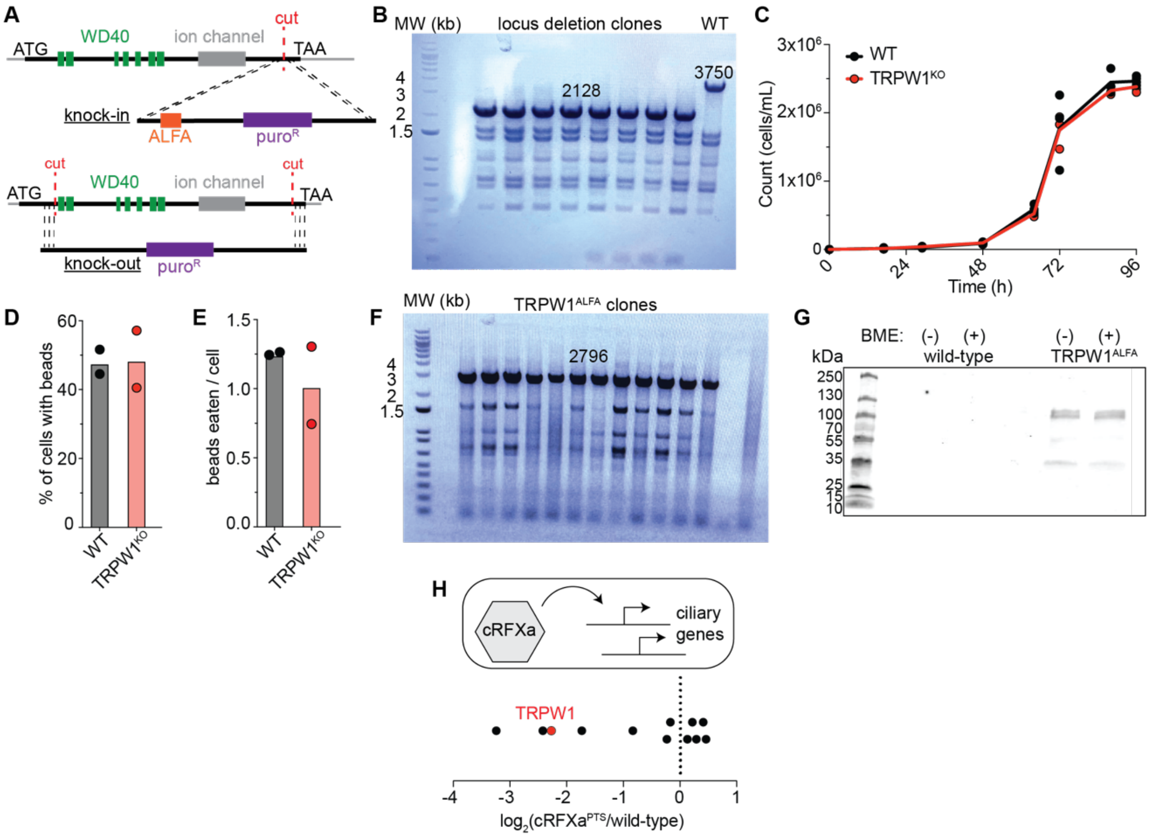


**Figure S3. Genome editing and genetic analysis of TRPW1.** (**A**) CRISPR-Cas9 genome editing strategy for TRPW1 alleles, derived from Booth *et. al.* (*19*) and Combredet *et. al* (*20*). Double-stranded DNA repair templates were generated by PCR from a plasmid encoding a puromycin resistance gene flanked by *S. rosetta* gene regulatory elements. See Table S3 for oligonucleotide sequences. (**B**) A DNA agarose gel showing successful locus deletion of TRPW1. Primers annealing on either side of the TRPW1 locus form a 3750 bp band in wild-type cells. In puromycin-resistant knockout clones, the expected size of 2128 is observed. (**C**) The TRPW1$^{KO}$ strain grows equivalently to wild-type *S. rosetta*. n = 3 technical replicates per time point, representative of n = 2 experiments. (**D**) TRPW1$^{KO}$ strain and wild-type cells show equivalent phagocytosis of 1 µm red fluorescent polystyrene beads, as quantified by the percent of cells eating beads. n = 2 technical replicates, representative of n = 3 experiments. (**E**) TRPW1$^{KO}$ strain and wild-type cells show equivalent phagocytosis of 1 µm red fluorescent polystyrene beads, as quantified by the number of ingested beads per cell. n = 2 technical replicates, representative of n = 3 experiments. (**F**) DNA agarose gel showing successful ALFA-tagging for the C-terminus of TRPW1. Primers flanking the C-terminus should yield a 2796 bp product if successful insertion of the ALFA tag, 3' UTR, and puromycin resistance cassette was achieved. (**G**) Lysates of TRPW1$^{ALFA}$ choanoflagellates show immunoreactivity with a nanobody against the ALFA tag, near the predicted molecular weight of 129 kDa for a TRPW1 monomer. (**H**) Multiple TRPW gene family members, including TRPW1, are down-regulated in a cRFXa mutant, showing co-regulation with a flagellar transcriptional program (*39*). Original data included n = 4 biological replicates for RNA sequencing. p-value < 0.0001 for down-regulation of TRPW1.

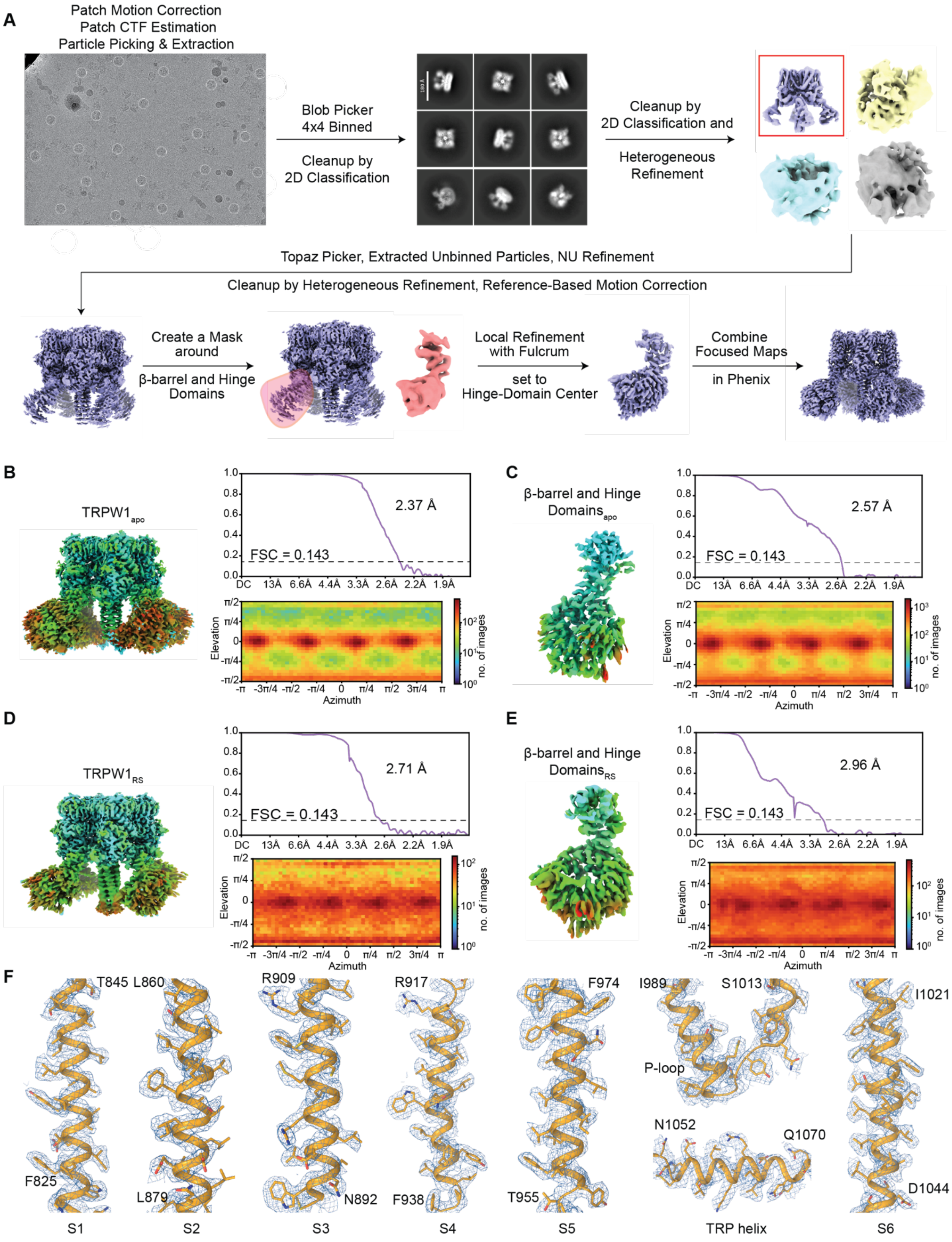

A
Patch Motion Correction
Patch CTF Estimation
Particle Picking & Extraction
Blob Picker
4x4 Binned
Cleanup by
2D Classification
Cleanup by
2D Classification and
Heterogeneous
Refinement
Topaz Picker, Extracted Unbinned Particles, NU Refinement
Cleanup by Heterogeneous Refinement, Reference-Based Motion Correction
Create a Mask
around
β-barrel and Hinge
Domains
Local Refinement
with Fulcrum
set to
Hinge-Domain Center
Combine
Focused Maps
in Phenix
B
TRPW1apo
2.37 Å
FSC = 0.143
C
β-barrel and Hinge
Domainsapo
2.57 Å
FSC = 0.143
D
TRPW1RS
2.71 Å
FSC = 0.143
E
β-barrel and Hinge
DomainsRS
2.96 Å
FSC = 0.143
Elevation
Azimuth
no. of images
F
T845 L860
R909
R917
F974
I989
S1013
I1021
P-loop
N1052
Q1070
F825
L879
N892
F938
T955
D1044
S1
S2
S3
S4
S5
TRP helix
S6

**Figure S4. cryo-EM analysis of TRPW1.** (**A**) Flowchart that outlines cryo-EM image acquisition and processing workflow performed to obtain the TRPW1 structures, with a representative micrograph showing example particles circled in white and 2D class averages. (**B-E**) Cryo-EM maps colored according to the local resolution estimation in cryoSPARC (left), FSC curves calculated between half maps, with the overall resolution estimated using the FSC = 0.143 criterion (top right) and angular distribution of particles calculated using the 3D refinement reconstruction algorithm in cryoSPARC (bottom right) for the consensus (B,D) and focused on β-barrel and hinge domain (C,E) reconstructions of TRPW1 in the apo (B-C) and RS 39604-bound (D-E) states. (**F**) Fragments of the apo-state TRPW1 TMD with the structural model shown as a ribbon and sticks and the corresponding cryo-EM density as a blue mesh.

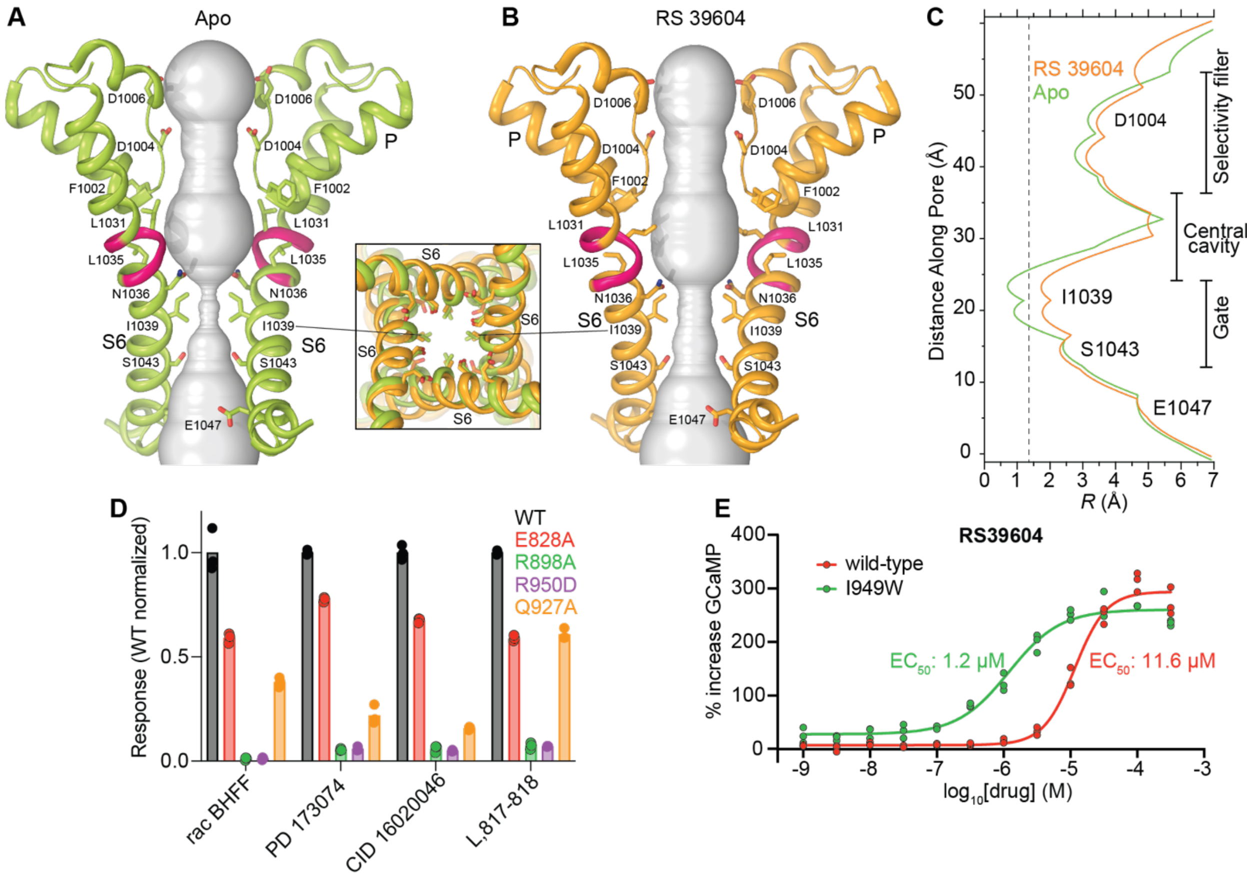


**Figure S5. Structure-function analysis of TRPW1.** (**A-B**) Pore-forming domain of TRPW1 in the apo (left, green) and RS 39604-bound (right, orange) states, with the residues contributing to the pore lining shown as sticks. Only two of four subunits are shown, with the front and back subunits omitted for clarity. The pore profiles are shown as space-filling models (grey). The π-bulge in S6 is highlighted (pink). The insert in the middle shows the intracellular view of the pore narrow constriction with the two structures superimposed. (**C**) Pore radius for apo (green) and RS 39604-bound (orange) structures of TRPW1 calculated using HOLE. The vertical dashed line denotes the radius of a water molecule, 1.4 Å. (**D**) The effect of single amino acid substitutions on the activation of heterologously expressed TRPW1 by the listed synthetic agonists (100 μM). Activity is shown normalized to the wild-type response. n = 3 $Ca^{2+}$ imaging replicates per drug. (**E**) A single amino acid substitution intended to improve the fit of RS 39604 in its binding pocket TRPW1 lowers the $EC_{50}$ approximately ten-fold. n = 3 $Ca^{2+}$ imaging replicates per concentration.

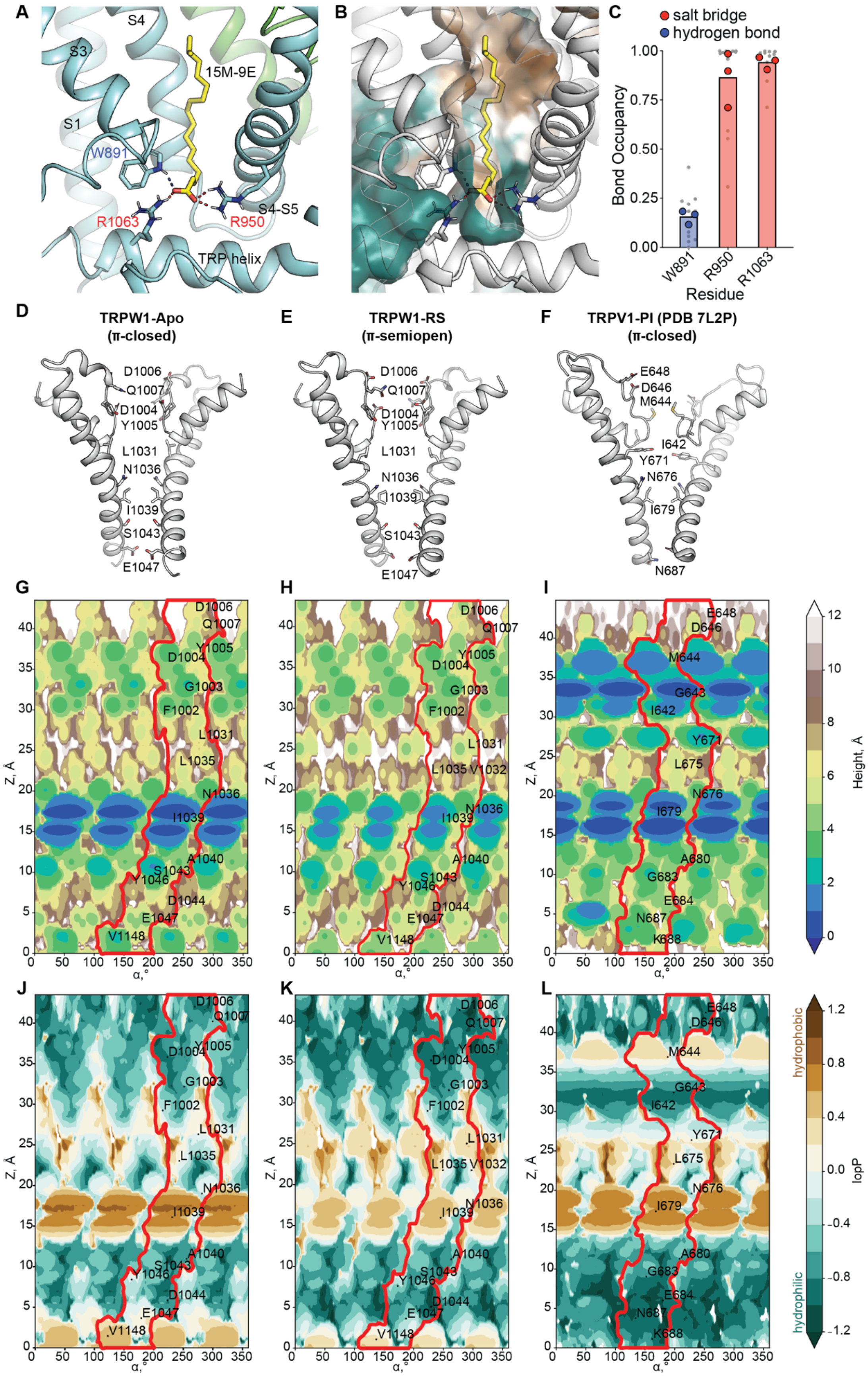
A
S4
S3
15M-9E
S1
W891
R1063
R950
S4-S5
TRP helix
B
C
salt bridge
hydrogen bond
Bond Occupancy
1.00
0.75
0.50
0.25
0.00
W891
R950
R1063
Residue
D
TRPW1-Apo
(π-closed)
D1006
Q1007
D1004
Y1005
L1031
N1036
I1039
S1043
E1047
E
TRPW1-RS
(π-semiopen)
F
TRPV1-PI (PDB 7L2P)
(π-closed)
E648
D646
M644
I642
Y671
N676
I679
N687
G
H
I
Z, Å
α,°
Height, Å
J
K
L
hydrophobic
hydrophilic
lopP
F1002
G1003
L1035
V1032
A1040
Y1046
D1044
V1148
G643
L675
A680
G683
E684
K688

**Figure S6. Molecular dynamics simulations of 15M-9E binding to TRPW1 and structural-physicochemical comparison of TRPW1 and TRPV1 ion-conducting pores.** (**A**) Representative MD snapshot of 15M-9E bound in the TRPW1 VBP. Residues forming salt bridges with the ligand are shown as sticks and highlighted in red; hydrogen-bonding residues are shown in blue. (**B**) Molecular hydrophobicity potential (MHP) mapped onto the surface of the VBP of TRPW1. Blue-green colors denote hydrophilic regions and brown denotes hydrophobic regions. The same state and viewing orientation are used as in panel A. (**C**) Average occupancy for salt bridges (red) and hydrogen bonds (blue) between 15M-9E and the TRPW1 vanilloid-binding site, obtained from n = 3 MD simulations. Gray dots are occupancy values for each tetramer subunit, larger colored dots are average across all 4 subunits during a simulation. (**D,E**) The ion-conducting pore of TRPW1 in the Apo- (D) and the RS 39604-bound (E) states. **(F)** The ion-conducting pore of TRPV1 in the PI-bound π-closed state (PDB 7L2P). In (D-F) the protein is shown in cartoon representation, and pore-lining residues are shown as sticks and labeled. (**G-I**) Cylindrical projections of the molecular hydrophobic potential (MHP) on the pore surface of TRPW1-Apo (G), TRPW1-RS 39604 (H) and TRPV1 (I). The vertical axis (Z) corresponds to the channel pore axis, and the horizontal axis indicates rotation around Z. Colors represent the distance from the pore axis to the pore surface, with the narrowest regions shown in blue. (**J-L**) Cylindrical projections of MHP on the pore surface of TRPW1-apo (J), TRPW1-RS 39604 (K) and TRPV1 (L). MHP values are expressed in octanol–water logP units (*36*). Projections were generated with the Molecular Surface Topography tool in the CELL framework (https://model.nmr.ru/cell). The area filled by one of the four subunits is marked by a red curve. Pore-lining residues are indicated on the projections. The ion-conducting pore of TRPW1 shows features conserved among the TRP-family channels: (i) a π-bulge segment in the central part of the pore and (ii) an activation gate formed by the nonpolar isoleucine side chains (I1039 for TRPW1 and I679 for TRPV1) and polar groups of asparagine residues (N1036 for TRPW1 and N676 for TRPV1, also see **Figure S5A,B**). These residues form a narrow constriction at the gate region (Z=15-20 Å), visible as blue in the landscape projections (G-I). In the closed TRPW1-apo (J) and TRPV1-PI (L) states, tightly packed isoleucine residues create a highly hydrophobic patch impermeable to water and ions, while the pre-activated TRPW1-RS39604 state shows markedly reduced hydrophobicity at the gate (K).

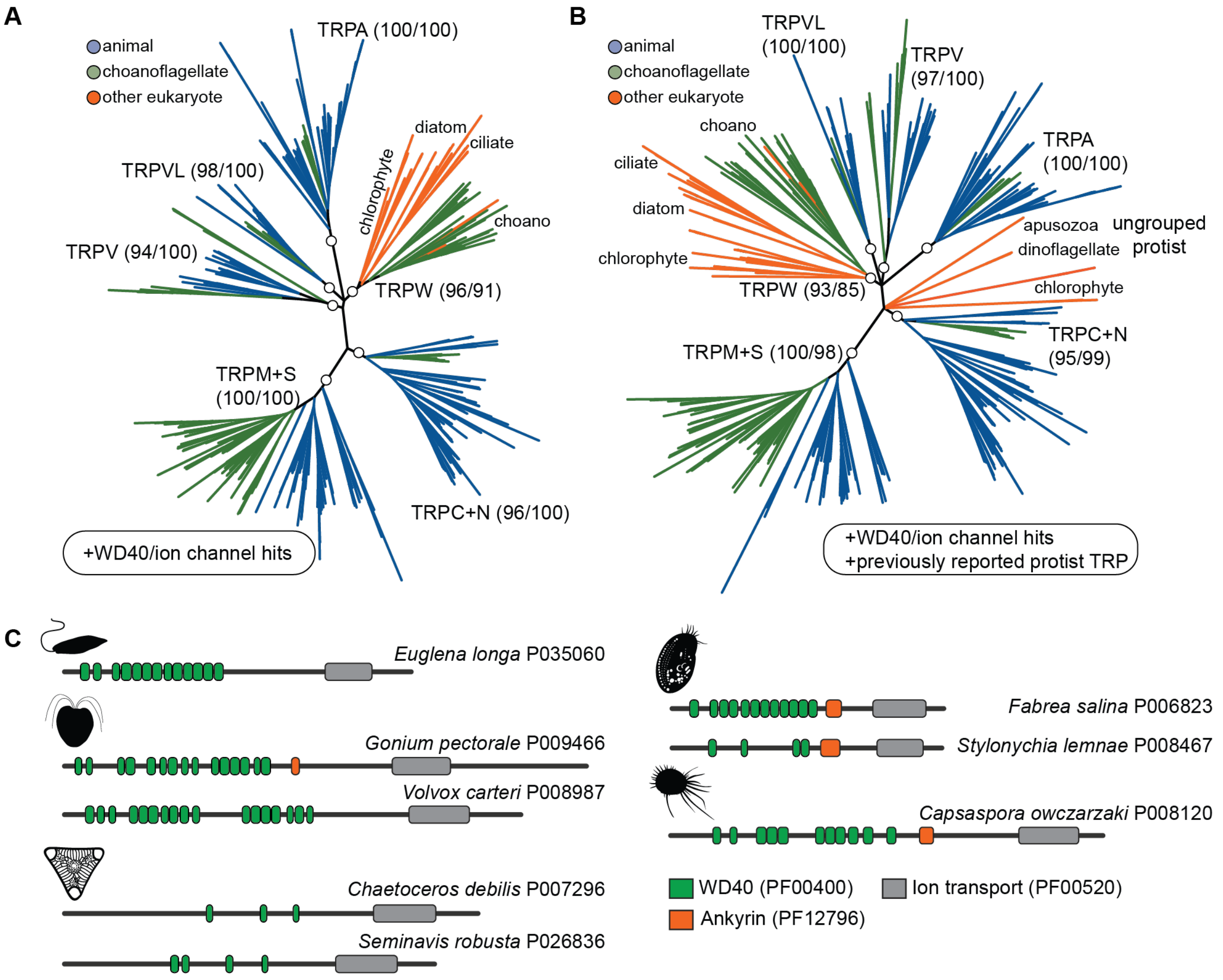


**Fig. S7. Evolutionary analyses for the TRPW family.** (**A**) The TRPW gene family contains homologs from other protists in addition to choanoflagellates. Diverse TRP channel sequences from choanoflagellates and animals were combined with genes detected in diverse eukaryotes by searching for the ion channel + WD40 repeat architecture. For display, branches with less than 70% bootstrap support by UFboot were collapsed into palintomies. Support values for circled nodes are listed as UFboot/SH-aLRT. The two orange sequences within the choano TRP channel clade are from *Euglena longa*. (**B**) Previously reported protist TRP channels group either with the TRPW family or do not group with any other family. "Ungrouped protist" sequences include reported TRP channels from chlorophyte algae (*15*), a reported TRPV from apusozoans (*11*), and two reported TRPMs from a dinoflagellate (*16*). For display, branches with less than 70% bootstrap support by UFboot were collapsed into palintomies. Support values for circled nodes are listed as UFboot/SH-aLRT. (**C**) Representative domain architectures from TRPW homologs in diverse eukaryotes.

#### Supplementary Movies

**Movie S1:** *S. rosetta* cells accumulate near a biofilm of *Algoriphagus machipongonensis* over a period of 1 hour, with frames taken every 10 seconds. Scale bar: 20 μm.

**Movie S2:** Unicellular *S. rosetta* exposed to the TRPW1 agonist 15M-9E switches stochastically between actively beating and paralyzed states. Scale bar: 5 μm.

**Movie S3:** Multicellular *S. rosetta* shows coordinated flagellar arrest when exposed to the TRPW1 agonist RS. Scale bar: 5 μm.

#### Supplementary Data Files

**Data S1:** FASTA file of predicted ion channel domains from *S. rosetta* genes with a hit for Pfam profile PF00520. Used for phylogenetic tree in Figure 1B. Sequences labeled by transcript accession number in *S. rosetta* genome as hosted on Ensembl protist.

**Data S2:** Results of searching the EukProt database for genes with the domain architecture of a voltage-gated superfamily transmembrane domain (PF00520) and WD40 domains (PF00400). File indicates the taxa with a hit, the number of genes with the architecture of interest, and the phylogenetic classification. Used for Figure 1C and Figure 5F. Taxa not represented in the phylogenetic tree in Figure 5F are indicated with asterisks. See Materials and Methods for discussion of single animal hits in *Pleurobrachia bachei* (ctenophore) and *Pristionchus pacificus* (nematode).

**Data S3:** FASTA file of predicted ion channel domains for TRP family sequences across animals and choanoflagellates, used for phylogenetic tree in Figure S1D. See Materials and Methods for details on how the sequence list was assembled. EukProt accession numbers provided.

**Data S4:** Differential expression of TRPW transcription in a cRFXa mutant compared to wild-type.

**Data S5:** FASTA file of ion channel domains from diverse animal and choanoflagellate TRP channels, as well as candidate TRPW orthologs from other protists. Used for phylogenetic tree in Figure S7A. See Materials and Methods for details on how the list of sequences was assembled.

**Data S6:** FASTA file including all sequences from Data S5, plus previously reported TRP sequences from protists. See Materials and Methods for details on how the list of sequences was assembled.

#### Supplementary Tables

**Table S1. Bacterial strains used for initial screening in this study**

| No. | Bacterial Strain | Strain ID |
|---|---|---|
| 1 | *Algoriphagus machipongonensis* | Amac |
| 2 | *Halomonas* sp. | Hsp |
| 3 | *Echinicola pacifica* | Epac |
| 4 | *Pseudomonas alcaligenes* | LP55 |
| 5 | *Pseudoalteromonas lipolytica* | LP30 |
| 6 | *Vibrio* sp. | LP27 |
| 7 | *Tenacibaculum litoreum* | LP39 |
| 8 | *Vibrio alginolyticus* | REJ23 |
| 9 | *Vibrio parahaemolyticus* | REJ3b |
| 10 | *Vibrio mytili* | OBE3 |
| 11 | *Halopseudomonas patchastrellae* | ORP22 |
| 12 | *Algoriphagus zhangzhouensis* | LP40 |
| 13 | *Vibrio mediterranei* | INC13 |
| 14 | *Tenacibaculum mesophilum* | LP44 |

**Table S2. $^1$H and $^{13}$C NMR chemical shift values of 15M-9E in $CD_3OD$**

| position | $\delta_C$[a] | type | $\delta_H$[b] | mult (*J* in Hz) |
|---|---|---|---|---|
| 1 | 180.1 | C | | |
| 2 | 36.9 | $CH_2$ | 2.21 | t (8.0) |
| 3 | 26.9 | $CH_2$ | 1.59 | m |
| 4–7 | 31.1–28.1 | $CH_2$ | 1.36–1.28 | m[c] |
| 8 | 27.9 | $CH_2$ | 2.04 | m |
| 9 | 130.6 | CH | 5.34 | m |
| 10 | 130.6 | CH | 5.34 | m |
| 11 | 27.9 | $CH_2$ | 2.04 | m |
| 12 | 30.6 | $CH_2$ | 1.32 | m[c] |
| 13 | 30.8 | $CH_2$ | 1.32 | m[c] |
| 14 | 40.1 | $CH_2$ | 1.18 | m |
| 15 | 28.9 | CH | 1.52 | m |
| 16 | 22.8 | $CH_3$ | 0.88 | d (6.5) |
| 17 | 22.8 | $CH_3$ | 0.88 | d (6.5) |

[a] 150 MHz, [b] 600 MHz, and [c] overlapped.

**Table S3. Oligonucleotide sequences for genome editing and genotyping.**

| Purpose | Sequence |
|---|---|
| Guide RNA near N-terminus of TRPW1 | TCCTGCGCGAGATCCATTGG |
| Guide RNA at C-terminus of TRPW1 | TGCAGAAGAAACAGTCTTTA |
| HDR template amplification, locus deletion, F | (ACGACGGCGGGTTGGAACTGCGCAACGTCTCATCTCCACCCACGCCGCCA)-TTTATTTAATTAAATAAA-GTAACGACTGCAGTCTTGTTGTC |
| HDR template amplification, locus deletion, R | (ACAGCACACAAAACAGCACACAAAACAGCACCATGCAGAAGAAACAGTC)-TTTTATTTAATTAAATAAA-CGGGCACCGCTCATTGATC |
| HDR template amplification, C-terminus ALFA tagging, F | (TCGAACAGCGCATCACGGCCCTCCTCGACCGCCTCCATCCCCGCCAGCCC)GGCTCTGGCTCTCGCC |
| HDR template amplification, C-terminus ALFA tagging, R | (ACAGCACACAAAACAGCACACAAAACAGCACCATGCAGAAGAAACAGTCT)GTAACGACTGCAGTCTTGTTGTC |
| Genotyping primer, C-terminus tagging, F | GTGGTTGCAGGGCATTTTCC |
| Genotyping primer, locus deletion, F | GAACTGCGCAACGTCTCATC |
| Genotyping primer for locus deletion and C-terminus tagging, R | ACACTTCCTCCTCTTCCCCC |

### Table S4. Cryo-EM data collection, refinement, and validation statistics

| **Structure** | **TRPW1$_{apo}$ Composite** | **TRPW1$_{apo}$ Consensus** | **TRPW1$_{apo}$ Beta-Barrel** | **TRPW1$_{RS}$ Composite** | **TRPW1$_{RS}$ Consensus** | **TRPW1$_{RS}$ Beta-Barrel** | **TRPW1$_{RS}$ Coiled-Coil** |
|---|---|---|---|---|---|---|---|
| EMDB code | EMD-77241 | EMD-77243 | EMD-77244 | EMD-77242 | EMD-77245 | EMD-77246 | EMD-77247 |
| PDB code | 35WH | - | - | 35WI | - | - | - |
| **Data collection and processing** | | | | | | | |
| Voltage (kV) | 300 | 300 | 300 | 300 | 300 | 300 | 300 |
| Electron exposure ($e^{-}Å^{-2}$) | 58 | 58 | 58 | 58 | 58 | 58 | 58 |
| Reported pixel size (Å) | 0.825 | 0.825 | 0.825 | 0.825 | 0.825 | 0.825 | 0.825 |
| Exposure number | 5,236 | 5,236 | 5,236 | 3,651 | 3,651 | 3,651 | 3,651 |
| **Processing software** | | | | | | | |
| Particle picking | cryoSPARC-v4.5.3 | cryoSPARC-v4.5.3 | cryoSPARC-v4.5.3 | cryoSPARC-v4.5.3 | cryoSPARC-v4.5.3 | cryoSPARC-v4.5.3 | cryoSPARC-v4.5.3 |
| Motion correction | Patch Motion Corr. | Patch Motion Corr. | Patch Motion Corr. | Patch Motion Corr. | Patch Motion Corr. | Patch Motion Corr. | Patch Motion Corr. |
| CTF estimation | Patch CTF | Patch CTF | Patch CTF | Patch CTF | Patch CTF | Patch CTF | Patch CTF |
| 2D/3D classification, refinement | cryoSPARC-v4.5 | cryoSPARC-v4.5 | cryoSPARC-v4.5 | cryoSPARC-v4.5 | cryoSPARC-v4.5 | cryoSPARC-v4.5 | cryoSPARC-v4.5 |
| Symmetry imposed | - | C4 | C1 | - | C4 | C1 | C1 |
| Initial particle number | - | 944,482 | 824,888 | - | 672,927 | 607,384 | 607,384 |
| Final particle number | - | 214,067 | 824,888 | - | 151,846 | 607,384 | 607,384 |
| Map resolution (Å) | - | 2.37 | 2.57 | - | 2.71 | 2.96 | 3.21 |
| **Refinement** | | | | | | | |
| FSC threshold | - | 0.143 | 0.143 | - | 0.143 | 0.143 | 0.143 |
| Map sharpening B factor ($Å^2$) | - | 60.1 | 74.4 | - | 70.6 | 63.8 | 138.2 |
| **Model composition** | | | | | | | |
| Non-hydrogen atoms | 25,776 | - | - | 31,021 | - | - | - |
| Protein residues | 3,344 | - | - | 3,332 | - | - | - |
| **Ligands** | | | | | | | |
| $Ca^{2+}$ | 4 | - | - | 4 | - | - | - |
| $Na^{+}$ | 0 | - | - | 1 | - | - | - |
| RS 39604 HCl | 0 | - | - | 4 | - | - | - |
| POV | 40 | - | - | 36 | - | - | - |
| Cholesterol | 4 | - | - | 4 | - | - | - |
| **B factors ($Å^2$)** | | | | | | | |
| Protein (mean) | 75.14 | - | - | 52.54 | - | - | - |
| Ligands (mean) | 21.84 | - | - | 45.53 | - | - | - |
| **R.m.s. deviations** | | | | | | | |
| Bond lengths (Å) | 0.00 | - | - | 0.00 | - | - | - |
| Bond angles (°) | 0.75 | - | - | 0.79 | - | - | - |
| **Validation** | | | | | | | |
| MolProbity score | 2.40 | - | - | 2.52 | - | - | - |
| Clash score | 3.27 | - | - | 6.07 | - | - | - |
| Poor rotamers (%) | 0.00 | - | - | 0.00 | - | - | - |
| **Ramachandran plot** | | | | | | | |
| Favored (%) | 89.98 | - | - | 90.88 | - | - | - |
| Allowed (%) | 9.75 | - | - | 9.00 | - | - | - |
| Disallowed (%) | 0.27 | - | - | 0.12 | - | - | - |

#### References


1. D. J. Richter, C. Berney, J. F. H. Strassert, Y.-P. Poh, E. K. Herman, S. A. Muñoz-Gómez, J. G. Wideman, F. Burki, C. de Vargas, EukProt: a database of genome-scale predicted proteins across the diversity of eukaryotes. *Peer Community Journal*, doi: 10.24072/pcjournal.173 (2022).
2. D. M. Emms, S. Kelly, OrthoFinder: phylogenetic orthology inference for comparative genomics. *Genome Biol.* **20**, 238 (2019).
3. J. Mistry, S. Chuguransky, L. Williams, M. Qureshi, G. A. Salazar, E. L. L. Sonnhammer, S. C. E. Tosatto, L. Paladin, S. Raj, L. J. Richardson, R. D. Finn, A. Bateman, Pfam: The protein families database in 2021. *Nucleic Acids Res.* **49**, D412–D419 (2021).
4. F. H. Yu, W. A. Catterall, The VGL-chanome: a protein superfamily specialized for electrical signaling and ionic homeostasis. *Sci. STKE* **2004**, re15 (2004).
5. S. R. Eddy, Accelerated profile HMM searches. *PLoS Comput. Biol.* **7**, e1002195 (2011).
6. S. F. Altschul, W. Gish, W. Miller, E. W. Myers, D. J. Lipman, Basic local alignment search tool. *J. Mol. Biol.* **215**, 403–410 (1990).
7. K. Katoh, D. M. Standley, MAFFT multiple sequence alignment software version 7: improvements in performance and usability. *Mol. Biol. Evol.* **30**, 772–780 (2013).
8. J. L. Steenwyk, T. J. Buida 3rd, Y. Li, X.-X. Shen, A. Rokas, ClipKIT: A multiple sequence alignment trimming software for accurate phylogenomic inference. *PLoS Biol.* **18**, e3001007 (2020).
9. L.-T. Nguyen, H. A. Schmidt, A. von Haeseler, B. Q. Minh, IQ-TREE: a fast and effective stochastic algorithm for estimating maximum-likelihood phylogenies. *Mol. Biol. Evol.* **32**, 268–274 (2015).
10. B. Q. Minh, M. A. T. Nguyen, A. von Haeseler, Ultrafast approximation for phylogenetic bootstrap. *Mol. Biol. Evol.* **30**, 1188–1195 (2013).
11. X. Cai, D. E. Clapham, Ancestral Ca2+ signaling machinery in early animal and fungal evolution. *Mol. Biol. Evol.* **29**, 91–100 (2012).
12. W. A. Valencia-Montoya, N. E. Pierce, N. W. Bellono, Evolution of sensory receptors. *Annu. Rev. Cell Dev. Biol.* **40**, 353–379 (2024).
13. N. J. Himmel, T. R. Gray, D. N. Cox, Phylogenetics identifies two eumetazoan TRPM clades and an eighth TRP family, TRP soromelastatin (TRPS). *Mol. Biol. Evol.* **37**, 2034–2044 (2020).
14. K. Fujiu, Y. Nakayama, H. Iida, M. Sokabe, K. Yoshimura, Mechanoreception in motile flagella of Chlamydomonas. *Nat. Cell Biol.* **13**, 630–632 (2011).
15. L. Arias-Darraz, D. Cabezas, C. K. Colenso, M. Alegría-Arcos, F. Bravo-Moraga, I. Varas-Concha, Brauchi, A transient receptor potential ion channel in Chlamydomonas shares key features with sensory transduction-associated TRP channels in mammals. *The Plant Cell* **27**, 177–188 (2015).

16. J. B. Lindström, N. T. Pierce, M. I. Latz, Role of TRP channels in dinoflagellate mechanotransduction. *Biol. Bull.* **233**, 151–167 (2017).

17. D. Oshima, M. Yoshida, K. Saga, N. Ito, M. Tsuji, A. Isu, N. Watanabe, K.-I. Wakabayashi, K. Yoshimura, Mechanoresponses mediated by the TRP11 channel in cilia of Chlamydomonas reinhardtii. *iScience* **26**, 107926 (2023).

18. A. Goehring, C.-H. Lee, K. H. Wang, J. C. Michel, D. P. Claxton, I. Baconguis, T. Althoff, S. Fischer, K. C. Garcia, E. Gouaux, Screening and large-scale expression of membrane proteins in mammalian cells for structural studies. *Nat. Protoc.* **9**, 2574–2585 (2014).

19. D. S. Booth, N. King, Genome editing enables reverse genetics of multicellular development in the choanoflagellate Salpingoeca rosetta. *Elife* **9**, e56193 (2020).

20. C. Combredet, M. Ansel, T. Brunet, A selection-based knockout approach for a choanoflagellate reveals regulation of multicellular development by Hippo signaling. *Cell Rep.* **44**, 116345 (2025).

21. M. H. T. Nguyen, I. S. Hernandez, F. U. Rutaganira, An accessible transfection protocol for choanoflagellates, *bioRxiv* (2026). https://doi.org/10.64898/2026.03.10.710884.

22. D. S. Booth, H. Szmidt-Middleton, N. King, Transfection of choanoflagellates illuminates their cell biology and the ancestry of animal septins. *Mol. Biol. Cell* **29**, 3026–3038 (2018).

23. C. Suloway, J. Pulokas, D. Fellmann, A. Cheng, F. Guerra, J. Quispe, Carragher, Automated molecular microscopy: the new Leginon system. *Journal of structural biology* **151**, 41–60 (2005).

24. A. Punjani, J. L. Rubinstein, D. J. Fleet, M. A. Brubaker, cryoSPARC: algorithms for rapid unsupervised cryo-EM structure determination. *Nat. Methods* **14**, 290–296 (2017).

25. E. F. Pettersen, T. D. Goddard, C. C. Huang, E. C. Meng, G. S. Couch, T. I. Croll, J. H. Morris, T. E. Ferrin, UCSF ChimeraX: Structure visualization for researchers, educators, and developers. *Protein Sci.* **30**, 70–82 (2021).

26. M. J. Abraham, T. Murtola, R. Schulz, S. Páll, J. C. Smith, B. Hess, E. Lindahl, GROMACS: High performance molecular simulations through multi-level parallelism from laptops to supercomputers. *SoftwareX* **1–2**, 19–25 (2015).

27. J. Huang, A. D. MacKerell Jr, CHARMM36 all-atom additive protein force field: validation based on comparison to NMR data. *J. Comput. Chem.* **34**, 2135–2145 (2013).

28. R. M. Venable, Y. Luo, K. Gawrisch, B. Roux, R. W. Pastor, Simulations of anionic lipid membranes: development of interaction-specific ion parameters and validation using NMR data. *J. Phys. Chem. B* **117**, 10183–10192 (2013).

29. W. L. Jorgensen, J. Tirado-Rives, Potential energy functions for atomic-level simulations of water and organic and biomolecular systems. *Proc. Natl. Acad. Sci. U. S. A.* **102**, 6665–6670 (2005).

30. B. Hess, H. Bekker, H. J. C. Berendsen, J. G. E. M. Fraaije, LINCS: A linear constraint solver for molecular simulations. *J. Comput. Chem.* **18**, 1463–1472 (1997).

31. G. Bussi, D. Donadio, M. Parrinello, Canonical sampling through velocity rescaling. *J. Chem. Phys.* **126**, 014101 (2007).

32. M. Parrinello, A. Rahman, Polymorphic transitions in single crystals: A new molecular dynamics method. *J. Appl. Phys.* **52**, 7182–7190 (1981).

33. T. Darden, D. York, L. Pedersen, Particle mesh Ewald: An N log (N) method for Ewald sums in large systems. *Journal of chemical physics* **98**, 10089–10092 (1993).

34. K. Vanommeslaeghe, E. Hatcher, C. Acharya, S. Kundu, S. Zhong, J. Shim, E. Darian, O. Guvench, P. Lopes, I. Vorobyov, A. D. Mackerell Jr, CHARMM general force field: A force field for drug-like molecules compatible with the CHARMM all-atom additive biological force fields. *J. Comput. Chem.* **31**, 671–690 (2010).

35. W. Yu, X. He, K. Vanommeslaeghe, A. D. MacKerell Jr, Extension of the CHARMM General Force Field to sulfonyl-containing compounds and its utility in biomolecular simulations. *J. Comput. Chem.* **33**, 2451–2468 (2012).

36. Y. A. Trofimov, N. A. Krylov, A. S. Minakov, K. D. Nadezhdin, A. Neuberger, A. I. Sobolevsky, R. G. Efremov, Dynamic molecular portraits of ion-conducting pores characterize functional states of TRPV channels. *Commun. Chem.* **7**, 119 (2024).

37. S. A. Wildman, G. M. Crippen, Validation of DAPPER for 3D QSAR: conformational search and chirality metric. *J. Chem. Inf. Comput. Sci.* **43**, 629–636 (2003).

38. W. L. DeLano, Pymol: An open-source molecular graphics tool. *CCP4 Newsl. protein crystallogr* (2002).

39. M. C. Coyle, A. M. Tajima, F. Leon, S. P. Choksi, A. Yang, S. Espinoza, T. R. Hughes, J. F. Reiter, D. S. Booth, N. King, An RFX transcription factor regulates ciliogenesis in the closest living relatives of animals. *Curr. Biol.*, doi: 10.1016/j.cub.2023.07.022 (2023).

40. F. Leon, J. M. Espinoza-Esparza, V. Deng, M. C. Coyle, S. Espinoza, D. S. Booth, Cell differentiation controls iron assimilation in the choanoflagellate Salpingoeca rosetta. *mSphere* **10**, e0091724 (2025).
